# Mammalian circadian clock proteins form dynamic interacting microbodies distinct from phase separation

**DOI:** 10.1101/2023.10.19.563153

**Authors:** Pancheng Xie, Xiaowen Xie, Congrong Ye, Kevin M. Dean, Isara Laothamatas, S K Tahajjul Taufique, Joseph Takahashi, Shin Yamazaki, Ying Xu, Yi Liu

## Abstract

Liquid-liquid phase separation (LLPS) underlies diverse biological processes. Because most LLPS studies were performed in vitro or in cells that overexpress protein, the physiological relevance of LLPS is unclear. PERIOD proteins are central mammalian circadian clock components and interact with other clock proteins in the core circadian negative feedback loop. Different core clock proteins were previously shown to form large complexes. Here we show that when transgene was stably expressed, PER2 formed nuclear phosphorylation-dependent LLPS condensates that recruited other clock proteins. Super-resolution microscopy of endogenous PER2, however, revealed formation of circadian-controlled, rapidly diffusing microbodies that were resistant to protein concentration changes, hexanediol treatment, and loss of phosphorylation, indicating that they are distinct from the LLPS condensates caused by overexpression. Surprisingly, only a small fraction of endogenous PER2 microbodies transiently interact with endogenous BMAL1 and CRY1, a conclusion that was confirmed in cells and in mice tissues, suggesting an enzyme-like mechanism in the circadian negative feedback process. Together, these results demonstrate that the dynamic interactions of core clock proteins is a key feature of mammalian circadian clock mechanism and the importance of examining endogenous proteins in LLPS and circadian studies.

## Introduction

The core mammalian circadian oscillators consist of autoregulatory transcription- and translation-based negative feedback loops. In the core mammalian circadian negative feedback loops, the heterodimeric CLOCK/BMAL1 complex acts as the positive element that activates transcription of clock genes by binding to E-boxes on gene promoters to rhythmically activate transcription (1–3). PERIOD proteins PER1 and PER2 and CRYPTOCHROME proteins CRY1 and CRY2 have been shown to interact with each other and function as the negative elements in the negative feedback loop by interacting and inhibiting the CLOCK/BMAL1 complex, leading to the closing the circadian negative feedback loop (2, 4–10). The functional circadian negative feedback loops lead to rhythmic gene expression and robust rhythmic PER abundance (2, 11). PER proteins associate stably with casein kinase 1 (CK1), and PER proteins are progressively phosphorylated after their synthesis and become hyperphosphorylated during the late night (11–13). PER phosphorylation plays critical roles in the mammalian circadian clock function by regulation of PER stability, its repressor activity, and its subcellular localization (12, 14–18). Due to their stable association, PER acts a scaffold for CK1 to promote phosphorylation of PER and of CLOCK (19, 20). The PER-dependent CLOCK phosphorylation leads to efficient removal of CLOCK-BMAL1 complexes from E-boxes on DNA. In addition, CRYs can repress CLOCK-BMAL1 activity on DNA independently of PER (21–24). These dual feedback mechanisms ensure robust circadian rhythmicity. We showed previously that the disruption of the PER-CK1 interaction by mutating residues of the interaction interface abolishes PER phosphorylation, impairs circadian negative feedback process and results in severe loss of circadian clock gene expression (19).

Like the *Drosophila* PERIOD protein (25), most human PER2 protein regions flanking the PAS domain are predicted to be intrinsically disordered (Figure 1A). Previous analyses of core circadian clock components showed that PER, CRY, CK1δ, BMAL1, and CLOCK proteins can be immunoprecipitated with each other and most of these proteins are present in a 1.9-MDa nuclear repressor complex (2, 9, 10, 26). Single-particle electron microscopy revealed that the nuclear complexes are ∼40 nm in diameter (26). In addition, isothermal titration calorimetry assays using recombinant proteins indicate that mCRY1-mPER2 affinity has a Kd in the lower nanomolar range in vitro and CRY and PER can form a stable complex in vitro (4, 5). On the other hand, glycerol gradient and chromatography-based approaches, however, suggest the presence of mostly discreet CLOCK-BMAL1 and PER-CRY-CK1δ complexes in mice liver extracts (27). In addition, the peaks of PER2 and CRY1 in these fractionation assays did not co-migrate in these fractionation assays (27). ChIP assays revealed that PER, CRY, BMAL1, and CLOCK were found to be rhythmically enriched on chromatin E-boxes (28). Imaging studies using PER2 knock-in reporter in mice and cell lines revealed that PER proteins are highly enriched in the nucleus (28–31). By expressing fluorescent fusion proteins via lentiviral transduction in U2OS/NIH/3T3 cells, studies showed that PER2 is mostly immobile in the nucleus and PER, CRY, BMAL1, and CLOCK proteins have high affinity for each other and exhibit very low nuclear diffusion coefficients (which indicate the ability to diffuse in the nucleus) with that of PER2 the lowest (28, 32). Furthermore, fluorescence recovery after photobleaching assay results suggest that most PER2 proteins are immobile in the nucleus of SCN cells (29, 31). These results are consistent with the proposal that most clock components associate with each other on chromatin (28, 32). However, lentiviral transduction normally resulted in clock protein overexpression albeit not as severe as transient transfection. How endogenous clock proteins interact and function remain unclear.

**Figure 1.**
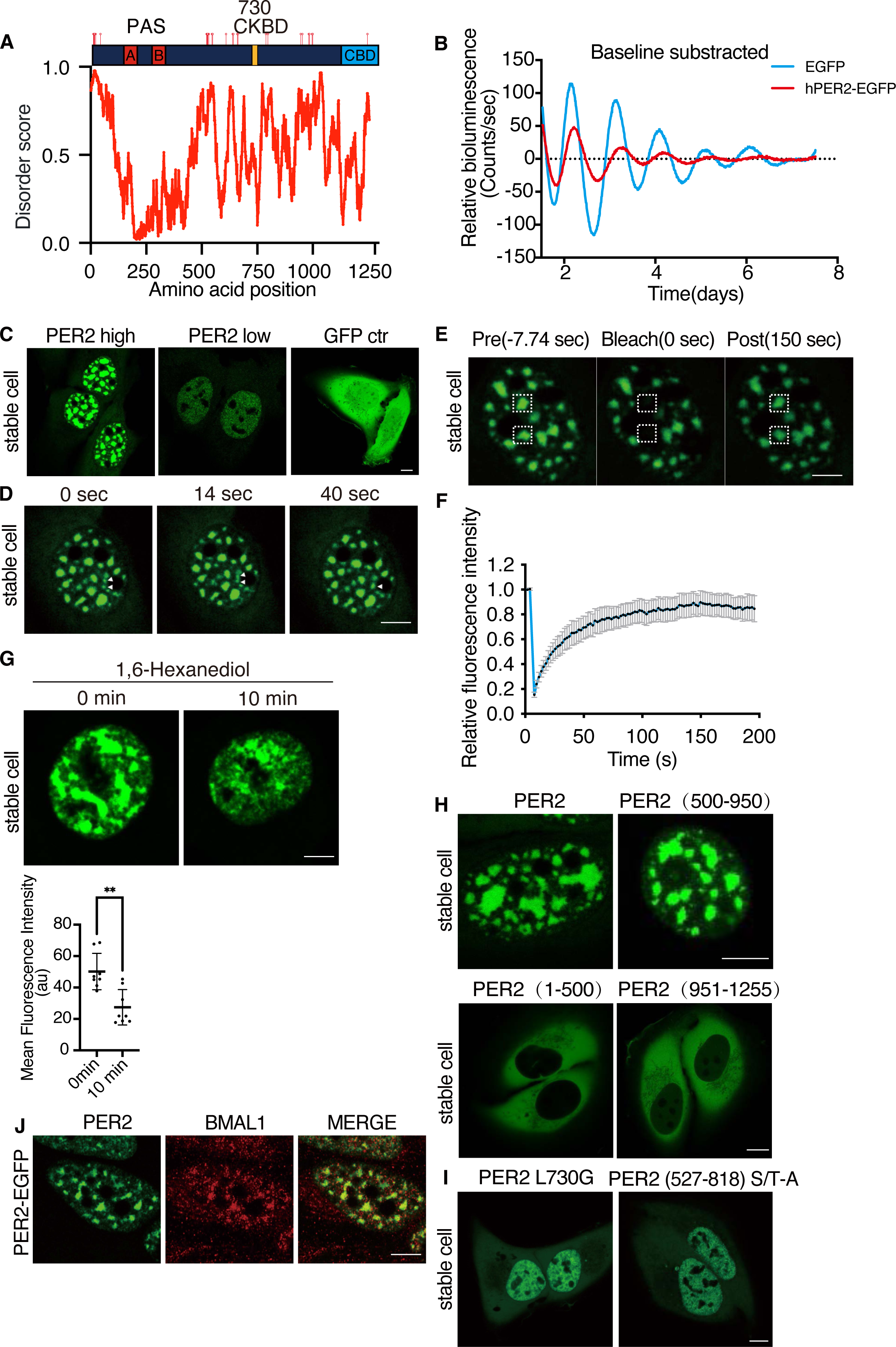
LLPS behavior of PER2-EGFP when it is constitutively overexpressed in U2OS cells. **(A)** Top: A diagram depicting human PER2 domains and phosphorylation sites. CKBD :Casein Kinase Binding domain. CBD: CRY binding domain. Location of the known phosphorylation sites were indicated based on a previously published data(12). Bottom: Disorder score across the human PER2 protein as predicted by IUPred. (**B)** Representative bioluminescence rhythms of the U2OS cells that stably express either EGFP or PER2-EGFP as well as an Per2(E2)-Luc reporter. Rhythms were monitored by LumiCycle and baseline-subtracted bioluminescence data are plotted (n=3 from three independent experiments). **(C)** Representative confocal images of cells stably express PER2-EGFP (high and low levels) and of control cells that express EGFP. Scale bar: 5 μm. (**D)** Images of a live cell that stably expresses PER2-EGFP over time. Two small condensates that fused during the time course are indicated with arrowheads. Scale bar: 5 μm. **(E)** Images of a live U2OS cell that stably expresses PER2-EGFP before and after photobleaching of regions marked by red squares. Scale bar: 5 μm. **(F)** Relative fluorescence intensity of PER2-EGFP condensates after photobleaching in the PER2-EGFP stable cells. Data are presented as mean ± SD (n=11). **(G)** Top: Images of live cells that express PER2-EGFP at 0 min and 10 min after treatment with 1,6-hexanediol (1.5%). Bottom: Quantification of mean fluorescence intensity of PER2-EGFP nuclei at 0 min and 10 min after treatment with 1,6-hexanediol. n= 8 cells. Data are presented as mean ± SD, unpaired two-tailed Student’s t test, ∗∗p < 0.01. **(H)** Representative confocal images of U2OS cells that overexpress PER2-EGFP, PER2(500-950)-EGFP, PER2(1-500)-EGFP or PER2(951-1255)-EGFP. PER2(1-500)-EGFP and PER2(951-1255)-EGFP protein are accumulated in the cytoplasm. (I) Representative confocal images cells that constitutively express PER2(L730G)-EGFP or PER2(527-818 S/T-A)-EGFP (PER2 with known and potential phosphorylation sites within aa 527-818 mutated to alanines). Scale bar: 5 μm. **(J)** Fluorescence immunostaining of endogenous BMAL1 in cells that stably overexpresses PER2-EGFP. Endogenous BMAL1 was stained with a BMAL1 antibody, and PER2-EGFP was monitored directly using the GFP channel. Scale bar: 5 μm.

Liquid-liquid phase separation (LLPS) results in formation of membraneless, spherical condensates of proteins or nucleic acids (33–35). An explosion of studies in the last decade have demonstrated that protein- and nucleic acid-containing LLPS condensates are involved in diverse biological processes including RNA metabolism, ribosome biogenesis, signal transduction, transcription, heterochromatin formation and immune response (33, 34, 36–39). The LLPS behavior of biomolecular molecules is concentration-dependent and is driven by multivalent macromolecular interactions. In many cases, intrinsically disorder regions (IDRs), also called low complexity domains (LCDs), have been shown to drive protein LLPS behavior due to their ability to mediate multivalent interactions (34, 40, 41). Characteristics of LLPS condensates include fusion of small condensates to become larger condensates and fluorescence recovery after photobleaching (33, 42, 43). Sensitivity of condensates to treatment of compounds such as 1,6-hexanediol, which interfere with weak multivalent hydrophobic interactions, is commonly used as an indicator of LLPS behavior (42–44). In addition, protein condensation is often regulated by post-translational modifications, and mutations that alter both LLPS and biological functions are also presumed to indicate physiological functions of LLPS (41, 45).

Although LLPS offers an attractive mechanism to explain many biological processes, most prior LLPS studies were based on protein overexpression in cells or in vitro biochemical studies (43). As a result, concerns have been raised regarding the physiological relevance of LLPS (42, 43). Nonetheless, the LLPS mechanism is often taken as physiologically relevant despite the lack of rigorous evidence on endogenous proteins. Careful examination of the herpes simplex virus replication compartments showed that despite characteristics of LLPS, these compartments operate via a mechanism different from LLPS. In addition, studies of RNA polymerase II and transcription factors led to the proposal that endogenous proteins may form “hubs” rather than LLPS condensates, which are observed when proteins are overexpressed (43, 46, 47). However, the same multivalent LCD-LCD interactions appear to mediate the formation of both protein hub and LLPS condensates.

Here we studied the dynamics and interactions of the endogenous core human circadian clock proteins. We observed that stably expressed PER2 from transgene forms nuclear condensates with typical LLPS-like behavior in a phosphorylation-dependent manner. Super-resolution fluorescence microscopy of the endogenous PER2, however, revealed that PER2 is present in rapidly diffusing nuclear microbodies that do not have LLPS characteristics and are resistant to loss of phosphorylation. These results demonstrate that the LLPS behavior of PER2 is caused by protein overexpression and may not be physiologically relevant. Although some endogenous PER2 do colocalize with BMAL1 and CRY1, surprisingly, most PER, BmaL1, and CRY1 exist in microbodies that are fast diffusing and do not colocalize in cells and in mice tissues. Together, these results demonstrate the dynamic nature of clock protein interactions, which requires revision of our current model of the mammalian clock mechanism. Together, our work highlights the importance of examining endogenous proteins in studies of the biological functions.

## Results

### LLPS behaviors of PER2-GFP when constitutively expressed in U2OS cells

To determine the cellular distribution of IDR-rich PER2 in human cells with a functional circadian clock, U2OS cells were transduced with a lentiviral vector for constitutive expression of a PER2-EGFP fusion protein under the control of the *EF-1a* promoter (Figure S1A). Control cells were transduced with the vector for expression of EGFP only. We did not use commonly used transient transfection to express the PER2-EGFP fusion protein because it resulted in approximately 100× higher levels of PER2-EGFP than detected in stably transduced cells (Figure S1B). We then transfected an Per2(E2)-Luc reporter plasmid into the U2OS cells that stably express PER2-EGFP and into control U2OS cells (48). The constitutive expression of PER-EGFP caused severe dampening of the circadian bioluminescence rhythm compared to control cells (Figure 1B). Constitutive overexpression of PER2 mediated by adenoviral infection was previously shown to abolish circadian rhythm in MEF cells (49). The dampened rhythm observed in our case is likely due to a reduced PER2-GFP overexpression level. Analysis using conventional confocal microscopy revealed that PER2-EGFP fluorescence was highly enriched in the nucleus (Figure 1C), which is similar to previous observations in the knock-in mouse cells and fibroblasts (29, 30). In contrast, upon transient transfection of a vector for expression of PER2-EGFP into U2OS cells, PER2-EGFP was mostly cytoplasmic during the first 2 to 3 days after transfection (Figure S1C), which is similar to previously reported (^50–52^). This difference is likely caused by the drastic overexpression of PER2-EGFP by transient transfection (Figure S1B). However, it is also important to note that the viral transduction resulted PER2-EGFP expression in stable cells still led to PER2 overexpression that is 20-fold higher than that of the endogenous PER2 level (see below).

After transduction, fluorescence signal was detected in about two dozen discrete puncta in the nucleus in most of the cells that express PER2-EGFP. These puncta were mainly spherical when small but large irregularly shaped condensates were also observed (Figure 1C). The small condensates were able to fuse, and this resulted in the larger one (Figure 1D and Supplemental data file 1). In cells with low levels of PER2-EGFP fluorescence, however, the PER2-EGFP condensates were not observed and signals were evenly distributed in the nucleus (Figure S1C), suggesting that the PER2-EGFP condensate formation is concentration-dependent. In contrast, in control cells with high EGFP expression level, fluorescence signal was evenly distributed in both cytoplasm and nucleus (Figure 1C), indicating that the condensate-behavior of PER2-EGFP is not caused by EGFP. Although EGFP is normally monomeric, it can dimerize at high concentrations (53). To exclude the effect of dimerization on our findings, we expressed the PER2 fused to EGFP(A206K); this mutation prevents dimerization (53). Condensate formation was similar whether PER2 was fused to EGFP or the mutant (Figure S1D). This result indicates that the condensation of PER2-EGFP is not caused by EGFP dimerization.

The PER2-EGFP condensates are somewhat stationary in the nuclei of live cells, which allowed us to perform the fluorescence recovery after photobleaching assay. EGFP fluorescence in the condensates showed a rapid recovery after photobleaching with a t_1/2_ of ∼23 s (Figure 1E-F). The alcohol 1,6-hexanediol interferes with weak hydrophobic interactions, and LLPS condensates dissolve upon treatment with 1,6-hexandiol (42, 43). When U2OS stable cells that express PER2-EGFP were treated with 1.5% 1,6-hexanediol, there was marked decrease of the fluorescence signal of the PER2-EGFP condensates after 10 min (Figure 1G) due to that monomeric PER2-eGFP protein is below the detection limit of the microscope. Together, these results demonstrate that constitutively expressed PER2 exhibits LLPS behavior in the nucleus.

LLPS behavior of many proteins is regulated by post-translational modifications and the correlations between protein mutations with biological effects and impacts on LLPS behavior have been used to support hypotheses that LLPS behavior is physiologically relevant (34, 42). PER phosphorylation has been shown to play important roles in circadian clock function (12, 13, 54). Thus, a role for PER phosphorylation in regulating its LLPS behavior would suggest that PER LLPS has a biological role. To identify the PER2 domain involved in LLPS, we created U2OS cell lines that stably express the N-terminal (amino acids 1-500), middle (amino acids 501-950), or C-terminal (amino acids 951-1255) regions of the protein fused to EGFP. We found that the expression of the middle region, but not the N- or C-terminal domains, resulted in nuclear PER2-EGFP condensates (Figure 1H). The PER2(1–500)-EGFP and PER2(951–1255)-EGFP proteins were found to be evenly distributed in the cytoplasm. The middle region of PER2 contains the previously identified PER-CK1 interaction domain and IDRs with many phosphorylation sites (Figure 1A) (12, 16, 19, 55). To determine the role of PER2 phosphorylation in condensate formation, we stably expressed PER2-EGFP with the L730G mutation in the PER-CK1 interaction domain, which disrupts the PER-CK1 interaction and abolishes PER2 phosphorylation in cells and in mice (19). Fluorescence in cells that expressed the mutant was distributed throughout the nucleus and no nuclear condensates were observed even though PER2(L730G)-EGFP was expressed at a much higher level than the PER2-EGFP (Figure 1I and S1E-F). The PER2-EGFP condensate formation was also abolished when 34 known and potential PER2 phosphorylation sites between aa 527-818 were mutated to alanine (Figure 1I and S1F). Thus, the formation of the PER2-EGFP condensates is dependent on PER phosphorylation.

To examine whether the PER2 condensates colocalize with the endogenous clock protein BMAL1, we performed immunofluorescence staining of the cells that stably express PER2-EGFP using a BMAL1-specific antibody. As shown in Figure 1J, most of the endogenous nuclear BMAl1 was found to localize in the PER2-GFP condensates, suggesting that the PER2 condensates recruit the BMAL1-CLOCK complex, which should sequester CLOCK-BMAL1 complex and inhibit its function in circadian transcriptional activation. Together, these results appear to support a model in which the phosphorylation-regulated LLPS properties of PER proteins are important for circadian clock function.

### Behavior of the endogenous PER2 tagged with EGFP by CRISPR/Cas9-mediated knock-in

To determine whether the endogenous PER2 exhibits LLPS behavior as does the ectopically expressed PER2, we modified a previously developed CRISPR/Cas9-based method (30) to tag the C-terminal end of the endogenous PER2 with EGFP reporter in U2OS cells (Figure 2A). Due to the low expression levels of the endogenous PER2, direct selection of correct homologous integration clones by EGFP fluorescence signal resulted in mostly false negative clones. In the donor vector, an mCherry reporter was added upstream of the *Per2* left homologous recombination arm and the sequence encoding the ECFP reporter was added between the EGFP reporter and the right homologous recombination arm (Figure 2A). Thus, fluorescence-activated cell sorting (FACS) of ECFP-positive but mCherry-negative cells from the blasticidin resistant cells could be used to greatly enrich for cells in which the donor vector had been integrated at the endogenous *Per2* genomic locus by double homologous recombination. Subsequently, the sequence encoding the ECFP reporter, which is flanked by LoxP sites, was removed by transfecting selected cells with a CRE expression plasmid; this restored the 3’UTR of the *Per2* locus.

**Figure 2.**
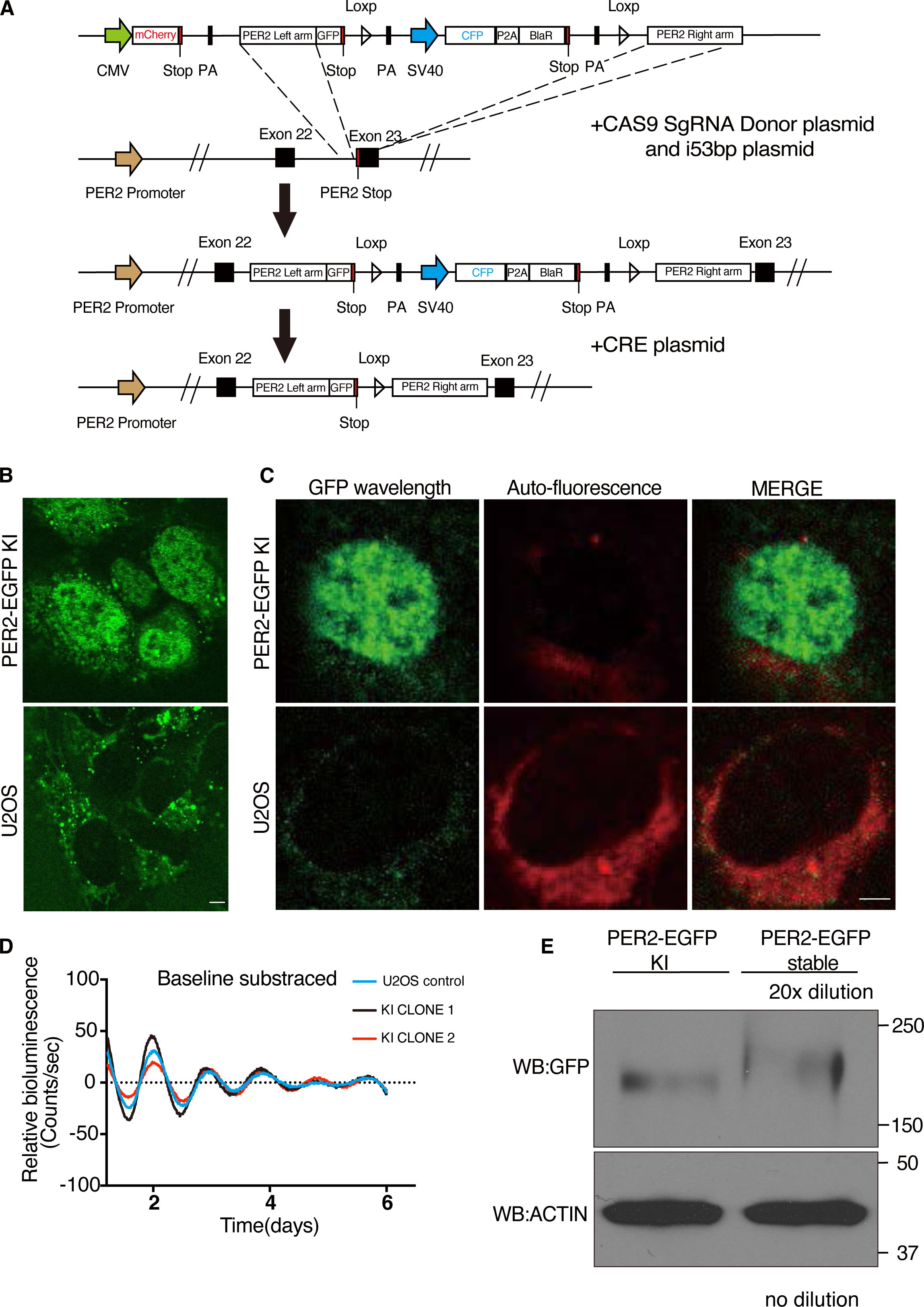
CRISPR/Cas9-mediated knock-in to tag the endogenous PER2 with EGFP. **(A)** Schematic of PER2-EGFP knock-in donor plasmid and genome editing strategy. (**B)** Live cell confocal imaging of PER2-EGFP KI U2OS cells and U2OS cells without KI. Scale bar: 5 μm. **(C)** Lambda scan of PER2-EGFP KI cells and control cells. The GFP wavelength was specifically extracted and all other wavelengths were merged as background. Scale bar: 5 μm. **(D)** Representative bioluminescence rhythms of the Per2(E2)-Luc reporter in the PER2-EGFP KI and control U2OS cells. Baseline-subtracted bioluminescence data are plotted (n=3). (**E)** Western blot showing the PER2-EGFP levels in the PER2-EGFP KI cells and cells that stably overexpress PER2-EGFP. The protein sample from the cells that stably overexpress PER2-EGFP was diluted 20 fold.

Single clones of the resulting ECFP-negative cells were obtained by FACS sorting. PCR using primer sets flanking the integration site followed by DNA sequencing confirmed the desired in-frame integration of *EGFP* gene before the stop codon of *Per2* in the knock-in (KI) cells. Examination by live cell fluorescence microscopy revealed that >40% of clones had highly enriched nuclear EGFP signal (Figure 2B), as previously reported for PER2-tagged reporter cells (29, 30). In contrast, there was no nuclear fluorescence signal in the control U2OS cells without the KI.

Fluorescence was also detected in the perinuclear regions in the cytoplasm of both knock-in and control U2OS cells (Figure 2B). Lambda scan of the fluorescent images was used to extract this background noise spectra and the EGFP signal. Most of the perinuclear signals were indeed background noise in the KI cells (Figure 2C). Due to the cytoplasmic background autofluorescence signals, we focused on the nuclear fluorescence signals. An Per2(E2)-Luc reporter plasmid was transfected into the PER2-EGFP KI and control U2OS cells (48). Our results revealed that both KI and control cells had similarly robust circadian bioluminescence rhythms with the same periodicity (Figure 2D). This indicated that the EGFP KI did not affect endogenous PER2 function, which is similar to previous reporter knock in at the PER2 C-terminus (31, 56).

Although cells that stably express PER2-EGFP also exhibited nuclei-enriched EGFP signals, comparison of PER2-EGFP protein levels by western blot analysis showed that the PER2-EGFP levels in cells that stably express this construct are more than 20-fold higher than the peak endogenous PER2-GFP levels (measured 10 h after dexamethasone synchronization) in the KI cells (Figure 2E). This result indicated that although the viral transduction-mediated PER2 expression in stable cells are much lower than that by transient transfection of cells, its levels are still much higher than those of the endogenous GFP-tagged PER2.

Consistent with previous analyses of the PER2 reporter KI cells (30), confocal florescence imaging of the live PER2-EGFP KI cells revealed a robust circadian rhythm of nuclear PER2-EGFP signal after the cells were synchronized by dexamethasone (DXMS) treatment. At its peak time (10 h after dexamethasone synchronization), the PER2-EGFP signal was found to be distributed throughout the nucleus; however, condensate-like structures were not observed in the KI cells (Figure 2B and 2C).

### Super-resolution imaging reveals circadian rhythm of the endogenous PER bodies

To determine the high-resolution spatial distribution of the endogenous PER2, we performed Airyscan super-resolution imaging of fixed PER2-EGFP KI cells at 10 h after DXMS synchronization (57, 58). Super-resolution imaging of fixed cells revealed hundreds of small PER2-EGFP foci in the nucleus (Figure 3A). To confirm that the nuclear EGFP foci contained PER2-EGFP, we also examined PER2-EGFP KI cells after treatment with *Per2*-specific siRNA and *Per2^KO^* cells (*Per2* gene was disrupted by CRISPR/Cas9 using a *Per2* specific sgRNA). The nuclear EGFP foci were abolished in cells depleted of PER2 (Figure 3B and S2A) but the perinuclear signals remained, further confirming that the perinuclear signals are due to autofluorescence. In live cells (scan time ∼ 1s), although most nuclear foci were blurred which should be due to their movements during the time needed for image collection, clear foci were still observed (Figure 3A and other live cell experiments below). In addition, the distribution of PER2-EGFP signals in live cells is very similar to previously shown for endogenous PER2 fusion with other fluorescent reporters in cells and mice (29–31). These results indicated that the small PER2-EGFP foci are the endogenous foci and not artifacts caused by cell fixation. These nuclear foci of PER2 were referred as PER bodies because their behavior and fluorescence intensities at the experimental condition used indicate that they are due to the presence of multiple PER2 molecules in each. Although some PER bodies were also detected in the cytoplasm, they were far fewer than in the nucleus. Moreover, signals from these bodies in the cytoplasm could be masked by the perinuclear autofluorescence signals. Thus, we focused our analyses on the nuclear PER bodies.

**Figure 3.**
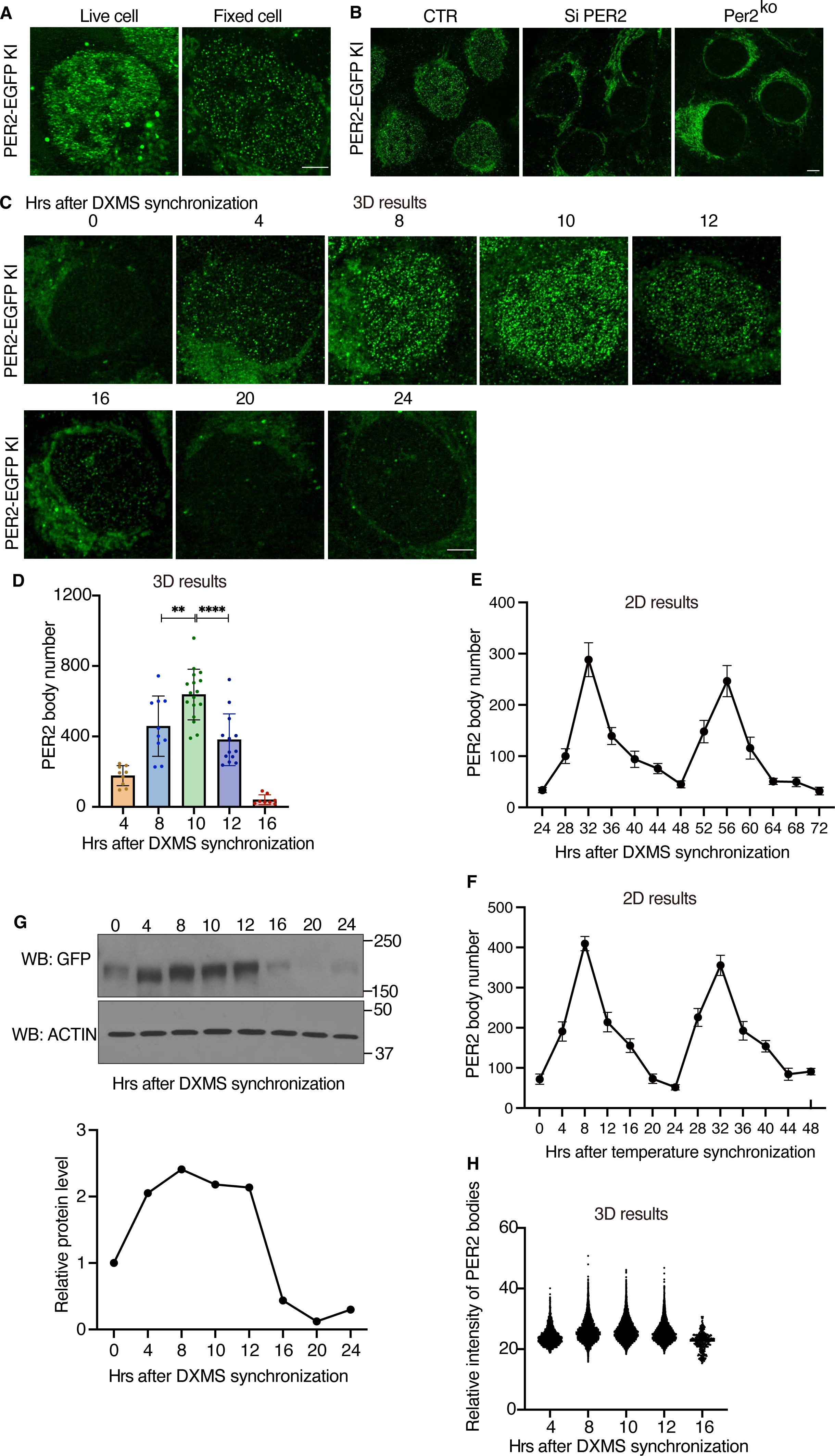
Super-resolution imaging of PER2-EGFP KI cells showing PER2 is concentrated in nuclear microbodies. **(A)** Airyscan confocal imaging of live (left) and fixed (right) PER2-EGFP KI cells in the nucleus. Scale bar: 5 μm. (**B)** Airyscan imaging of fixed PER2-EGFP KI cells (left), PER2-EGFP KI cells treated with *Per2*-specific siRNA (si-PER2), and *Per2*^KO^ cells (*Per2* gene disrupted by CRSPR/Cas9). Scale bar: 5 μm. (**C)** Time course 3D imaging results of nuclear GFP signals in PER2-EGFP KI cells after dexamethasone synchronization. Representative 2D images from the 3D results are shown. (**D)** Numbers of nuclear PER bodies at different time points after dexamethasone synchronization (n=8-17 cells). Data are presented as mean ± SD, n = (8-17) cells, unpaired two-tailed Student’s t test, ^∗∗^p < 0.01; ^∗∗∗∗^p < 0.0001. **(E)** Quantification of the PER2-EGFP microbodies of time course 2D imaging results of the PER2-EGFP KI cells after dexamethasone synchronization from hr 24-72. (**F**) Quantification of PER2-EGFP microbodies of time course 2D imaging results of of the PER2-EGFP KI cells after temperature cycle synchronization from hr 0-48. (**G)** Top: Western blot of EGFP in PER2-EGFP KI cells at indicated time points after dexamethasone synchronization. Equal amount of protein was loaded in each lane. Bottom: Quantification of PER2-EGFP levels over time of the western blot result. (**H)** Nuclear PER body mean fluorescence intensities at indicated time points after dexamethasone synchronization (n=8-17 cells).

To determine the number of nuclear PER bodies in a cell, we performed 3D Airyscan super-resolution microscopy of fixed PER2-EGFP KI cells harvested at different time points after cell synchronization by DXMS. The number of PER bodies exhibited a robust rhythm, with a peak at 10 h after synchronization at approximately 650 PER bodies per nucleus and a trough at 20-22 h after synchronization with approximately dozens of bodies per cell (Figure 3C-D, and Supplemental data file 2). In addition, robust rhythms of PER bodies with a similar phase and period as seen for the 0-24 hrs were seen 24-72 hrs after cell synchronization by DXMS and 0-48 hrs after temperature cycle entrainment of cells (Figure 3E-F and S2B-C), indicating that the rhythms of PER bodies in the first 24 hrs are genuine circadian rhythms and are not DXMS-induced artifact. In addition to the PER body rhythm, the PER2-EGFP protein levels also exhibited a robust rhythm with a peak at approximately 8 h and a trough at approximately 20 h after synchronization (Figure 3G). Comparison of these two rhythms revealed that although PER2-EGFP levels at 4 h were comparable to those at 10 h, the number of PER bodies was more than 3 times higher at hr 10 h than that at hr 4. In addition, the peak of PER bodies occurred at ∼10 h, whereas the PER2-EGFP levels peaked at 8 h. Nuclear and cytoplasmic fractions of the cell extracts showed that the nuclear PER2-EGFP level is only modestly higher at hr 10 than that at hr 4 (Figure S2D). Thus, PER level alone does not determine the formation of PER bodies and at hr 4, there are relatively more PER molecules are not in PER bodies and cannot be detected by microscope.

Although the fluorescence intensities of the nuclear PER bodies within each nucleus varied, their sizes did not appear to change significantly, indicating that the sizes of PER bodies are smaller than the resolution (∼140 nm) of the microscope. Importantly, although PER levels at 16 h were about 80% lower than those from at 8-12 h and PER body numbers were also much lower, fluorescence intensities of the PER bodies, which should reflect their sizes, were not markedly different (Figure 3H). This result contrasts with the concentration-dependent LLPS behavior exhibited by the PER2-EGFP condensates in cells that stably overexpress the fusion protein (Figure 1C), indicating that the formation of endogenous PER2-EGFP bodies is less sensitive to protein concentration changes than the LLPS behavior of the overexpressed PER2-EGFP. The robust rhythms of PER2-EGFP protein and fluorescence are similar to the published rhythms of PER2 (10, 59), indicating that EGFP-fusion does not have a significant impact on PER2 function and stability.

We used two different methods to estimate the number of PER2 molecules in each PER body. Mass spectrometry-based analyses previously determined that the PER2 levels peaked at about 13,000 copies per cell (60). Based on the number of nuclear PER bodies at the peak, we estimated that each PER body might have on average ∼20 PER2 molecules. We also used known amount of recombinant EGFP protein to estimate the absolute abundance of PER2-EGFP molecules per PER2-EGFP KI cell at 10 h after synchronization (Figure S2E). Using this method, we estimate that there are on average ∼23 PER2-EGFP molecules in each PER body. Thus, both methods yielded similar values. It should be noted that both methods likely overestimate the number of PER2 molecules in a PER body because not all PER2 molecules are in nuclear PER2 (cytoplasmic PER2 and nuclear PER2 existed in smaller protein complexes that could not be detected). The PER bodies should also expect to include PER1, PER2, and PER3 proteins given previously established interactions of these proteins (19, 26).

### Most PER bodies freely diffuse in the nucleus but some are immobile for less than 1 s

PER, CRY, BMAL1, and CLOCK proteins were thought to exist primarily in a 1.9-MDa nuclear repressor protein complex (26). Fluorescence correlation spectroscopy of PER2, BMAL1, CLOCK, and CRY1 proteins fused with fluorescent reporters overexpressed in cells have very low nuclear diffusion coefficients; that of PER2 is the lowest (<0.2 μm^2^/s), suggesting that these overexpressed proteins are mostly immobile due to their association with chromatin (28, 61). To examine spatial dynamics of the endogenous nuclear PER bodies, we performed live cell imaging of the PER2-EGFP KI cells using an Airyscan super-resolution microscope in 0.35-1 s sampling intervals (Supplemental data file 3). Shorter intervals are not possible due to low signal levels and photobleaching. Surprisingly, no stationary PER bodies (i.e., a body immobile in at least three consecutive frames) were detected, and it was not possible to track the movement of individual PER body due to its rapid movements. This result indicate that the endogenous PER bodies are distinct from the stationary condensates seen in cells that stably overexpress PER2-EGFP. Thus, endogenous PER bodies move rapidly in the nucleus and normally do not stay on chromatin for more than 1 s.

To determine the dynamics of nuclear PER body movements, we then examined live KI cells using a high-resolution laser-scanning oblique plane microscope (light sheet microscopy) (62, 63), which allows for high-spatiotemporal imaging of fluorescent molecules under gentle, non-phototoxic illumination. Even here, with an exposure time of 30 ms per plane, PER body dynamics were too fast to image in a volumetric format. Thus, we settled on imaging a single oblique cross-section of the nucleus at 30-ms intervals (Figure 4A and Supplemental data file 4). Tracking the trajectories of individual fluorescent PER bodies confirmed that vast majority were not stationary (Figure 4B, representative tracks 1, 2, 3, 4), indicating that most PER bodies are not associated with chromatin. Analysis of mean square displacement of the PER bodies as a function of time showed that they have a mean diffusion coefficient of 12.12±0.5117 μm^2^/s (Figure 4C-D), which is similar to free diffusion behavior previously shown for transcription factors (64). However, 0.6% trackable traces of PER bodies revealed that some PER bodies were immobile for longer than 90 ms (Figure 4B). In one example (track 5), the PER body was immobile during the entire trackable duration, and in another (track 6), the body first moved and then became immobile. The average time of immobility of the immobile PER2 bodies was ∼0.4 s (Figure 4E). Our estimate of the percentage of PER bodies that are immobile is certainly an underestimate since we could not detect PER bodies that were immobile for less than 90 ms and one PER body can be responsible for multiple tracks due to the imaging limitation of single nuclear cross-section. Nonetheless, these results demonstrate that only a very small fraction of PER bodies are on chromatin at any given time and that PER bodies can only remain on chromatin for less than 1 s. The free diffusion behavior for most endogenous PER bodies is in contrast with previous fluorescence correlation spectroscopy analyses of cells that overexpress PER2 fusion protein (28). Our data indicate that lentivirus-based transduction results in PER2 expression levels much higher than that of the endogenous PER2, which causes the observed LLPS-like behavior.

**Figure 4.**
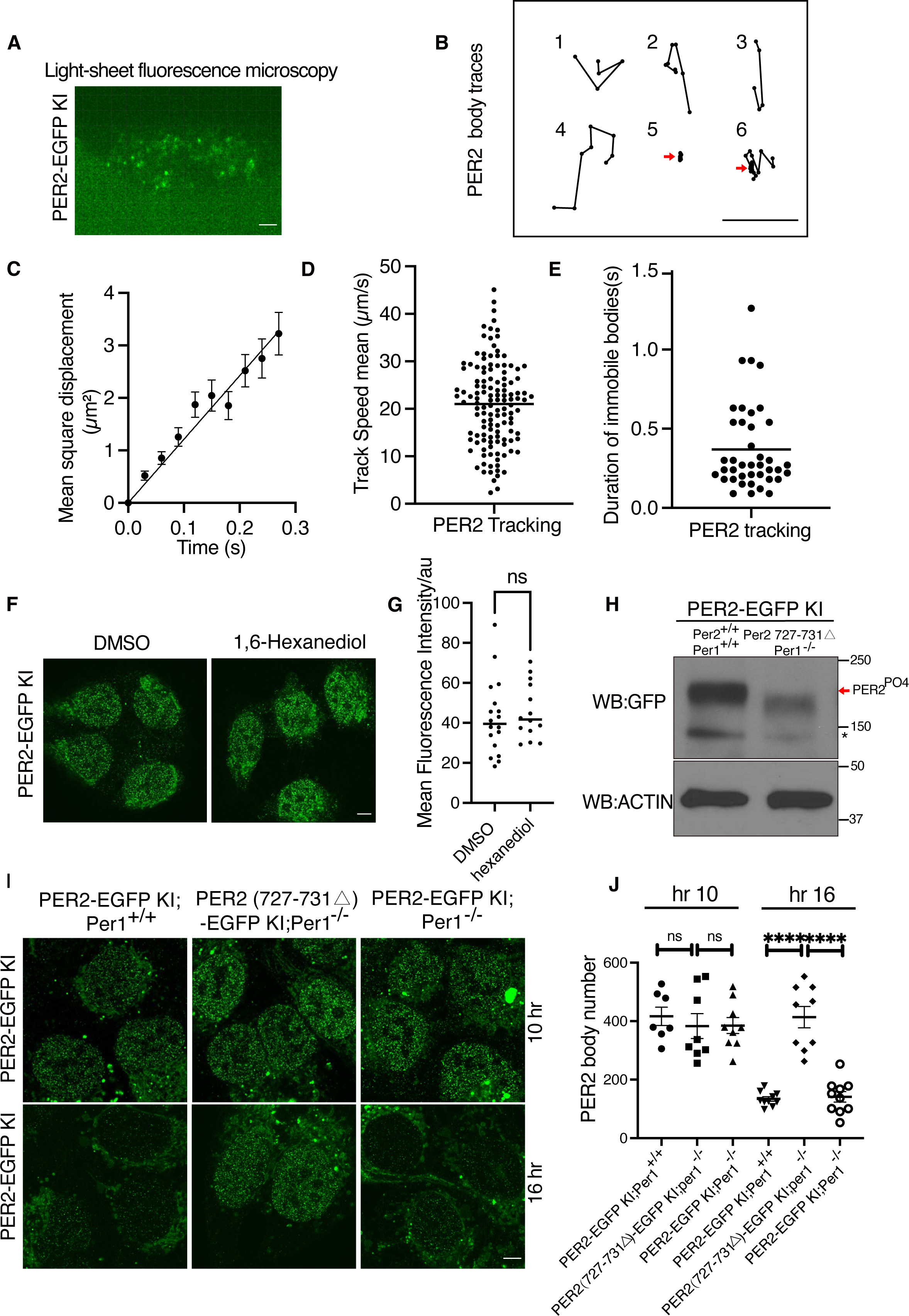
Most PER2 bodies diffuse freely in nucleus and are formed by mechanism distinct from overexpression-caused LLPS condensates. **(A)** Representative image of nuclear PER bodies in PER2-EGFP KI cell under light-sheet fluorescence microscopy at hr 10 after synchronization. Scale bar: 5 μm. (**B)** Representative movement tracks of PER bodies. Tracks 1-4: freely diffusing tracks. Tracks 5: immobile tracks. Track 6: a diffusing PER body became immobile. Scale bar: 5 µm. (**C)** The averaged mean square displacement (MSD) of PER bodies computed from data collected from 3 cells. Data are presented as means ± SEM. (**D)** Mean speeds of different tracks (n=3 cells). (**E)** Duration of immobility of PER bodies of the identified immobile PER bodies (data collected from 3 cells). (**F)** Representative image of nuclear PER bodies at 0 min and 10 min after treatment with 1,6-hexanediol in fixed KI cells at hr 10 after synchronization. Scale bar: 5 μm. (**G)** Comparison of nuclear PER2-EGFP fluorescence intensities of KI cells between 0 min and 10 min after treatment with 1,6-hexanediol at hr 10 after synchronization. Data are presented as median fluorescence intensity in each cells. n=14-18 cells. (**H)** Western blot of PER2-EGFP in the PER2-EGFP KI and the PER2(Δ727-731)-EGFP KI; *Per1*^KO^ cells at hr 10 after synchronization. The arrow indicates the phosphorylated PER2-EGFP species. * indicates a background protein band. (**I)** Representative Airyscan images of PER bodies in fixed PER2-EGFP KI and PER2(Δ727-731)-EGFP KI; *Per1*^KO^ cells at hr 10 after synchronization. Scale bar: 5 μm. (**J)** Nuclear PER2 body numbers in the PER2-EGFP KI, PER2-EGFP KI; *Per1*^KO^, and PER2(Δ727-73)-EGFP KI; *Per1*^KO^ cells at 10 h and 16 h after dexamethasone synchronization.

### PER bodies and LLPS condensates are formed by distinct mechanisms

It was previously proposed that the multivalent protein-protein interactions mediated by IDRs/LCDs of some transcription factors can mediate the formation of protein “hubs” at physiological concentrations (43, 46, 65). When these proteins are overexpressed, the same LCD-LCD multivalent interactions can result in formation of LLPS condensates (46). To determine whether the endogenous PER bodies are mediated by the same protein-protein interactions that result in PER2-EGFP-containing LLPS condensates, we treated with PER2-EGFP knock-in cells with 1, 6-hexanediol (1.5%), which rapidly abolished the condensates in cells that stably overexpressed PER2-EGFP (Figure 1G). In contrast, the hexanediol treatment did not affect number or fluorescence intensities of the PER bodies in the KI cells (Figure 4F-G and S3A). This result indicates that the endogenous PER bodies and the LLPS condensates are formed by distinct mechanisms: LLPS condensates are mediated by multivalent weak interactions (likely involving PER IDRs), whereas the endogenous PER bodies are formed by stronger protein-protein interactions that are not disrupted by hexanediol.

That LLPS behavior was abolished by mutation of the PER2-CK1 interaction domain or by mutation of PER2 phosphorylation sites (Figure 1H-I) prompted us to examine whether the endogenous PER bodies are sensitive to PER2 phosphorylation. We used the CRISPR/Cas9 method to mutate the PER2-CK1 interaction domain of the *Per2* gene in the KI cells. We obtained a single clone-derived cell line in which *Per2* has an in frame-deletion of five amino acids (Δ727-731) in the PER2-CK1 interaction domain (19). To exclude the effect of PER1 in mediating CK1-dependent PER2 phosphorylation, we further disrupted the endogenous *Per1* gene by CRSPR/Cas9 in these cells. As expected, the PER2-EGFP protein in the PER2(Δ727-731)-EGFP; *Per1*^KO^ cells became severely hypophosphorylated compared to the parental cells (Figure 4H). Despite the loss of PER2 phosphorylation, the numbers and intensities of the PER bodies were high in the PER2(Δ727-731)-EGFP KI; *Per1*^KO^ cells at 10 h after synchronization (Figure 4I-J and S3B-C). Whereas the number of PER bodies decreased dramatically at 16 h after synchronization in both PER2-EGFP KI and PER2-EGFP KI; *Per1*^KO^ cells, their numbers and their intensities remained high in the PER2(Δ727-731)-EGFP KI; *Per1*^KO^ cells. This result is consistent with our previous study showing the importance of the PER-CK1 interaction for clock function at the molecular level (19). These experiments also demonstrated that formation of endogenous PER bodies is not dependent on PER2 phosphorylation and the PER-CK1 interaction, further confirming that PER body formation and LLPS of PER2 are mediated by distinct mechanisms. Due to heterogeneity of PER2 microbodies, we cannot exclude the possibility that a few of small LLPS-dependent structures can still exist in cells. Thus, the PER2 LLPS condensates are mostly if not all caused by protein overexpression and are not physiologically relevant.

### Interactions among PER2-, BMAL1-, and CRY1-containing microbodies are dynamic

In cells that stably overexpress PER2-EGFP, most BMAL1 was recruited to the PER2-EGFP condensates (Figure 1J). Similarly, the overexpression of EGFP-PER2 was previously shown to dramatically reduce the mobility of CRY1 (28). These protein overexpression-based results are consistent with the proposal that most core clock proteins, including PER, are in large nuclear complexes on the chromatin (26). To examine the spatial distributions of other endogenous core clock proteins and their interactions, we inserted an in-frame mScarlet-I tag at the C-terminal end of the open reading frame of the endogenous *Bmal1* and *Cry1* loci in the PER2-EGFP KI cells using the CRSPR/Cas9 approach (Figure 5A). The resulting clonal cells express PER2-EGFP and BMAL1 or CRY1 specifically labeled with mScarlet-I at the C-terminus (Figure S4A). After synchronization by DXMS, cells were harvested at different time points, fixed to collect 2D images using Airyscan fluorescence microscopy. As expected, PER2-EGFP fluorescence exhibited a robust rhythm that peaked at 8-10 h after synchronization (Figure 5B-C). In contrast, the rhythm of BMAL1-mScarlet-I fluorescence was ∼ 8 hr phase advanced from the PER2 rhythm: BMAL1-mScarlet-I fluorescence peaked at 4 h and troughed at 10-12 h (Figure 5B), consistent with the advanced BMAL1 protein phase in mice (19, 66). On the other hand, despite the robust PER2-EGFP fluorescence rhythm, the CRY1-mScarlet-I fluorescence in the CRY1-mScarlet-I fluorescence did not exhibit a robust rhythm in the double KI cells (Figure 5C). Like PER bodies, hundreds of BMAL1-mScarlet-I and CRY1-mScarlet-I foci were also detected in each cell after fixation in the 2D imaging (Figure 5B-C), suggesting that BMAL1 and CRY1 also form multimeric complexes. Importantly, the merged images showed that most of the PER bodies did not colocalize with BMAL1-mScarlet-I/CRY1-mScarlet-I foci (Figure S4B-C). Similar results were also seen for cells at hr 32 after synchronization (S4D). Live cell imaging of these double KI cells further confirmed that the distribution patterns of the PER, BMAL1 and CRY1 bodies are very similar to those of the fixed cells (Figure 5D-E), further indicating that the results of the fixed cells are not artifacts caused by cell fixation procedure. Due to the rapid movements of these protein bodies and the time required for imaging in two separate channels, live imaging cannot be used for colocalization studies. In live cells, we also seen some large round fluorescent foci near to the edge of nuclear membrane, which are due to some unknown autofluorescent (revealed by Lambda scan) cellular structures.

**Figure 5.**
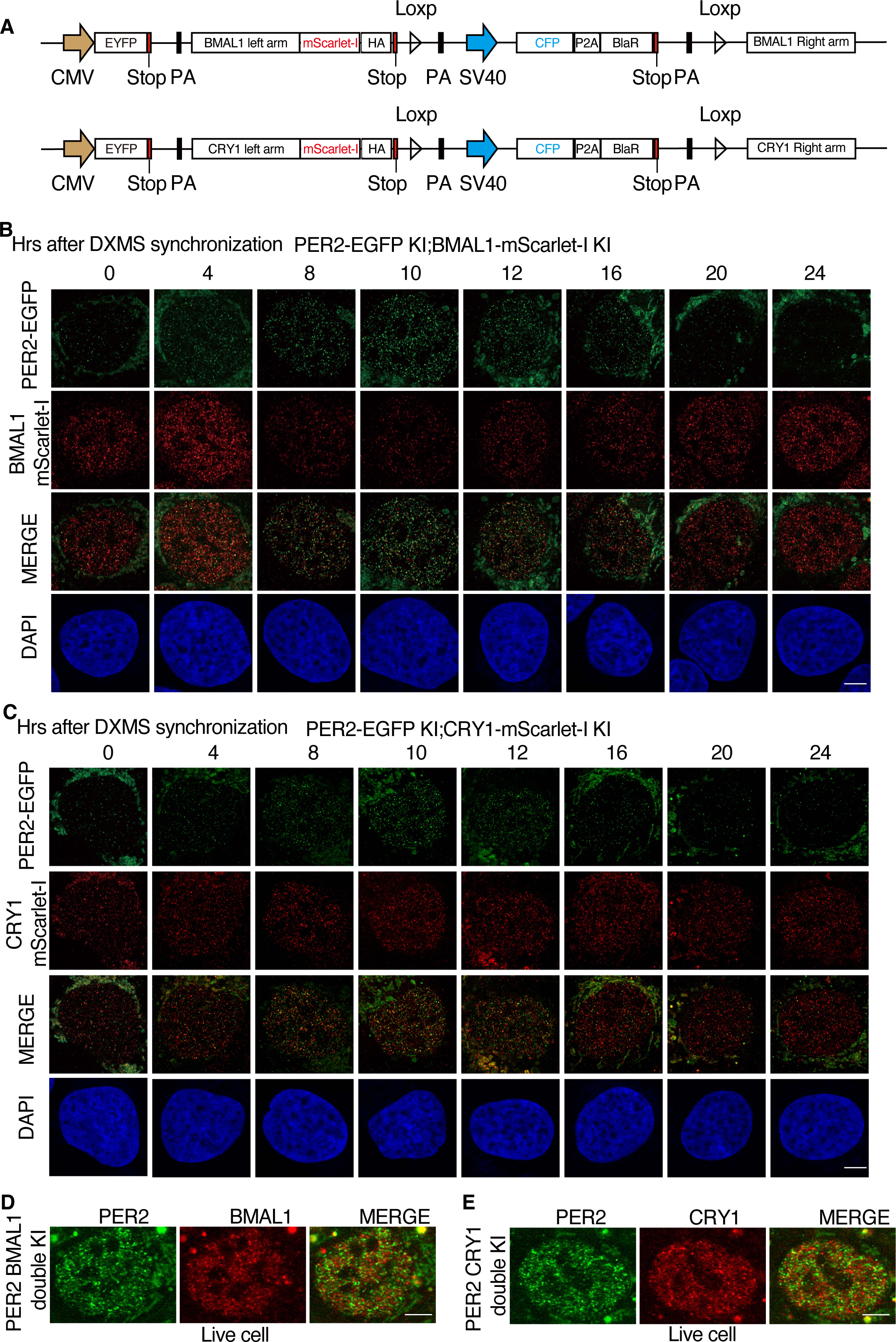
Airyscan imaging analyses of dual fluorescence reporter U2OS cells in which the endogenous PER2 and BMAL/CRY1 are tagged by EGFP and mScarlet-I, respectively. **(A)** Diagrams showing the design of BMAL1-mScarlet-I and CRY1-mScarlet-I knock-in donor plasmids. (**B)** Time course Airyscan fluorescence imaging of PER2 and BMAL1 in fixed PER2-EGFP and BMAL1-mScarlet-I double KI cells after dexamethasone synchronization. Scale bar: 5 μm. (**C)** Time course Airyscan imaging of PER2 and CRY1 in fixed PER2-EGFP and CRY1-mScarlet-I double KI cells after dexamethasone synchronization. Scale bar: 5 μm. (**D**) Airyscan fluorescence imaging of PER2 and BMAL1 in live PER2-EGFP and BMAL1-mScarlet-I double KI cells. (**E**) Airyscan fluorescence imaging of PER2 and BMAL1 in live fixed PER2-EGFP and CRY1-mScarlet-I double KI cells.

To estimate the numbers of BMAL1/CRY1 microbodies and identify the colocalization foci with PER2 bodies, we performed 3D Airyscan fluorescence microscopy analyses (Figure 6A-B and Supplemental data file 5-6), which revealed that the numbers of BMAL1 bodies were rhythmic but with a much lower amplitude than that of PER body rhythm (Figure 6C). There were about 800 BMAL1 bodies at its peaks (hr 4), and at troughs, there were about 5-600 BMAL1 foci. On the other hand, the number of CRY1 bodies stayed ∼600 at different time points after hr 0. Based on the previous estimates of ∼20,000 BMAL1 and CRY1 molecules per cell (60), there are likely ∼25-30 BMAL1 and CRY1 molecules in their respective bodies if all BMAL1 and CRY1 reside in their respective microbodies.

**Figure 6.**
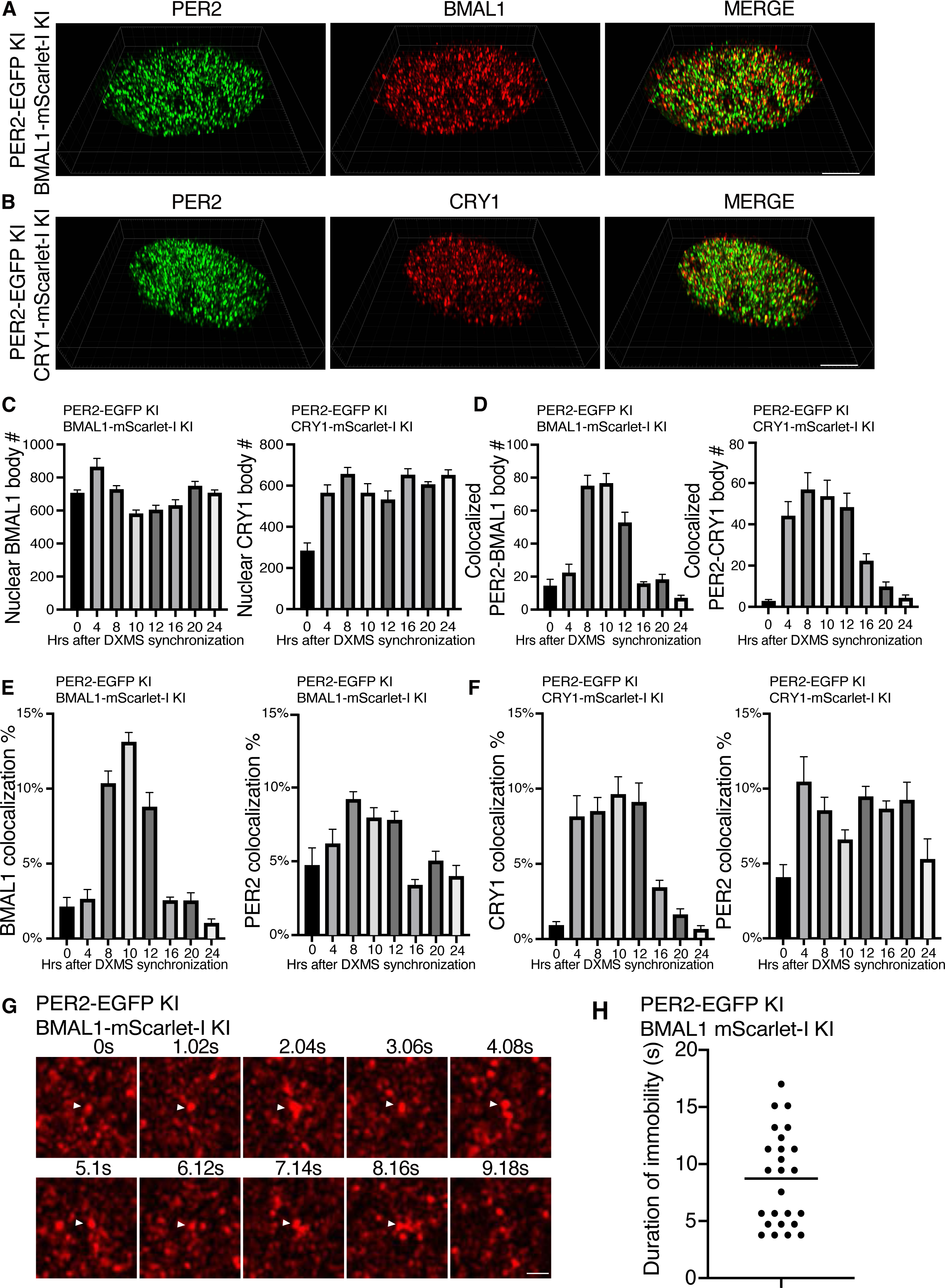
Airyscan imaging analyses of double KI cells showing that most PER2 bodies do not associate with BMAL1/CRY1 and the DNA residence time of BMAL1. **(A)** 3D Airyscan imaging result of PER2-EGFP and BMAL1-mScarlet-I double KI cells at hr 10 after synchronization. Representative converted 2D nuclear images were shown. The 3D Airyscan imaging results for other time points are presented in the supplemental data files. Scale bar: 5 μm. (**B)** 3D imaging of PER2-EGFP and CRY1-mScarlet-I double KI cells at hr 10 after synchronization. Representative converted 2D nuclear images were shown. Scale bar: 5 μm. (**C)** Numbers of BMAL1 bodies and CRY1 bodies at different time points after dexamethasone synchronization in the double KI cells. Data are means ± SEM (n=8 cells). (**D)** Numbers of PER2 and BMAL/CRY1 foci that colocalize as a function of time after dexamethasone synchronization. Threshold for colocalization was set < 100 nm. Data are presented as means ± SE (n=8 cells). (**E)** Percentages of BMAL1 and PER2 foci that colocalize at different time points after synchronization in the PER2 and BMAL1 double KI cells. Data are presented as means ± SEM (n=8 cells). (**F)** Percentage of CRY1 and PER2 foci that colocalize at different time points after synchronization in the PER2 and CRY1 double KI cells. Data are presented as means ± SEM (n=8 cells). (**G)** Time series of live cell Airyscan imaging of BMAL1 foci in the BMAL1-mScarlet-I knock-in cells. A selected nuclear region was shown. The arrow indicate an immobile BMAL1 foci. Frame rate: 1.02 s/frame. Scale bar: 5 μm. (**H)** Summary of duration of immobility of identified immobile BMAL1 foci in the BMAL1-mScarlet-I KI cells. Data are presented as median (n=10 cells).

To examine whether these proteins colocalize, we analyzed the 3D fluorescence images of the double-labeled cells and identified EGFP foci (PER2) and mScarlet-I (BMAL1/CRY1) foci that were less than 100 nm apart based on their centers using Imaris spots colocalization analysis (Figure S5A-B), which were regarded as interacting with each other if the sizes of PER2/BMAL1/CRY1 bodies are 100nm or larger. Most EGFP (PER2) and mScarlet-I (BMAL1 and CRY1) foci did not colocalize (Figure 6A-B and S5A-B). There were rhythms of the number of colocalized foci peaking at 8-10 h post synchronization for both PER2 with BMAL1 and PER2 with CRY1 (Figure 6D), corresponding to the time of peak PER body number. In addition, the percentages of PER2, BMAL1 and CRY1 bodies that colocalize with each other were also rhythmic (Figure 6E-F). At the peak of PER2 level after synchronization (hr 8-10), there were less than 10% of PER2 were considered to interact with BMAL1 and CRY1, respectively. For BMAL1 and CRY1, peak percentages associated with PER2 peaked at about 13% and 10%, respectively. The peaks of PER2 colocalization with BMAL1/CRY1 correlate with peak of PER2 levels, which is consistent with previously immunoprecipitation results (10). It is important to note that these colocalization results should be overestimates as PER, BMAL1, and CRY1 bodies that are smaller than the 100-nm colocalization threshold used (see below). Single-particle electron microscopy previously estimated the super nuclear PER-containing complexes from mouse liver to be ∼40-nm structures (26). When colocalization distance of 40 nm was used in our analysis, only about 2% of PER bodies were found to be colocalized with BMAL1 or CRY1 at h 10 (Figure S6A-C). Therefore, vast majority of PER2 molecules are not in complex with BMAL1 or CRY1 in the nucleus at any given time, indicating that their interactions are transient.

### Nuclear movements of BMAL1 and CRY1 bodies

We next examined nuclear dynamics of BMAL1-mScarlet-I and CRY1-mScarlet-I in the KI cells by performing live cell Airyscan super-resolution imaging at 1-s intervals (Supplemental data file 7-8). The low signal levels of the mScarlet-I-labeled proteins in the knock-in cells prevented us from examining the live cell imaging of BMAL1 and CRY1 bodies in shorter time intervals. BMAL1 and CRY1 bodies that stayed in the same location for three consecutive frames (∼3 s) were considered as immobile. We were not able to identify any CRY1 bodies that were immobile using this method, suggesting that CRY1 most freely diffuse and can only stay on chromatin for less than 3 s. For BMAL1, however, some immobile bodies were identified with many immobile for 3 to 5 s or as long as 16 s (Figure 6G-H), consistent with the residence time of known transcription factors on chromatin (64). Figure 6G show a BMAL1 body was immobile for 8 s before disappearing afterwards. These results indicate that BMAL1/CLOCK are on the chromatin considerably longer than PER and CRY proteins. The short duration of PER and CRY association with BMAL1/CLOCK on chromatin suggest that the PER and CRY bodies are enzyme-like and can act on multiple BMAL1/CLOCK complex, thus, resulting in efficient inhibition of the BMAL1-CLOCK complex activity and the removal of BMAL1-CLOCK complex from chromatin.

### Spatial distributions of PER2, BMAL1, and CRY1 and their associations revealed by STED microscopy and immunodepletion assays

Stimulated emission depletion (STED) fluorescence microscopy can circumvent the optical diffraction limit and can achieve lateral spatial resolution of ∼40 nm in cells (67, 68). We therefore used 2D STED immunofluorescence microscopy to determine the nuclear distribution of PER2-EGFP, BMAL1, and CRY1 proteins in the PER2-EGFP KI cells at 10 h after synchronization. We screened commercially available GFP, BMAL1, and CRY1 antibodies that can specifically detect PER2-EGFP, BMAL1, and CRY1 in the PER2-EGFP KI cells but not in control U2OS cells, MEF cells lacking BMAL1 or U2OS depleted for CRY1, respectively (59). As shown in Figure S7A-C, Anti-GFP, anti-BMAL1, and anti-CRY1 antibodies detected nuclear specific PER2-EGFP, BMAL1, and CRY1 foci in the PER2-EGFP KI cells but their signals were either absent or dramatically reduced in the control cells, indicating the specificity of these antibodies. 2D STED microscopy using two different antibodies (anti-GFP and anti-BMAL1 or anti-GFP and anti-CRY1) performed on the PER2-EGFP KI cells at 10 h after synchronization revealed hundreds of nuclear PER2-EGFP, BMAL1, and CRY1 foci per cell, consist with our Airyscan fluorescence data (Figure 7A-B). Merged images revealed that the vast majority of PER2-EGFP and BMAL1foci and of PER2-EGFP and CRY1 foci were not colocalized. Analysis of these 2D STED images revealed that 15-20% of the PER2-EGFP were less than 100 nm from centers of BMAL1 or CRY1 foci (Figure 7C-D). However, it should be noted that compared to 3D results, 2D analyses typically result in overestimation of protein colocalization by several folds.

**Figure 7.**
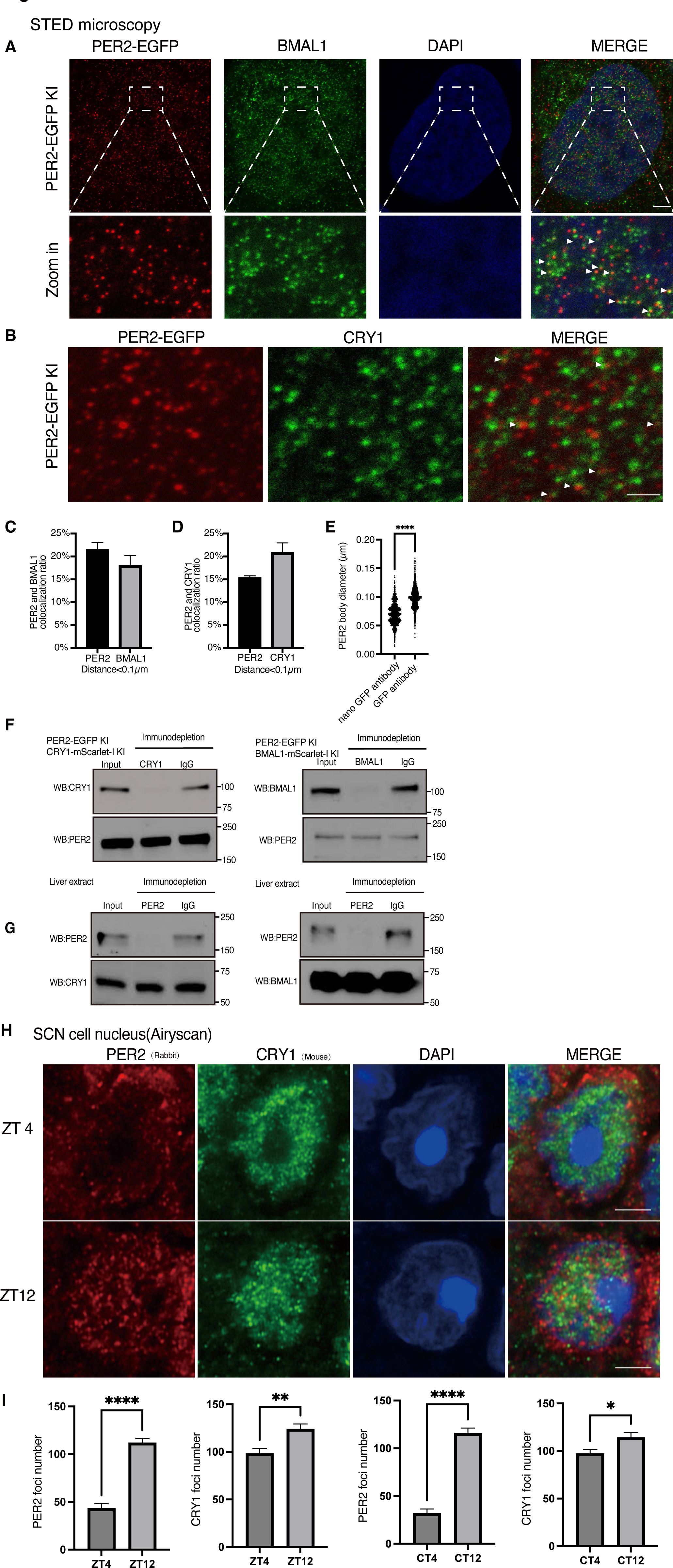
PER, BMAL1 and CRY1 microbodies and their interactions revealed by STED microscopy. **(A)** STED microscopy images of PER2 and BMAL1 puncta in the PER2-EGFP KI cells at 10 h after synchronization. PER2 and BMAL1 were stained by anti-GFP (Mouse) and anti-BMAL1 antibody (Rabbit), respectively. Scale bar: 5 μm. **B:** STED microscopy images of PER2 and CRY1 foci in the PER2-EGFP KI cells at 10 h after synchronization. PER2 and CRY1 were stained by anti-GFP and anti-CRY1 antibody, respectively. Scale bar: 5 μm. (**C)** Percentages of PER2 and BMAL1 bodies that colocalize at 10 h after synchronization. Distance threshold was set < 100 nm. Data are presented as mean ± SEM (n=8 cells). (**D)** Percentages of PER2 and CRY1 bodies that colocalize at 10 h after synchronization. Distance threshold was set < 100nm. Data are presented as mean ± SEM (n=3 cells). (**E)** Comparison of PER body sizes estimated by conventional anti-GFP antibody and anti-GFP nanobody. Data are presented as mean ± SEM (n=5 cells). (**F**) Immunodepletion assay of CRY1 or BMAL1 in extracts prepared from the indicated double KI U2OS cells 10 h after synchronization. After immunodepletion, western blot analyses were performed using the indicated antibodies. (**G**) Immunodepletion assays of mPER2 in extracts prepared from mice liver tissue (ZT18). (**H**) Representative immunofluorescence imaging results of mPER2 and mCRY1 in the nucleus using mice SCN tissues harvested at the indicated ZT time points using a rabbit polyclonal PER2 antibody and a mouse CRY1 antibody, respectively. Cells from the core region of SCN were selected here. Scale bar: 5 μm. (I) Quantification of the nuclear PER2/CRY1 foci numbers in the SCN cells from tissues harvested at the indicated time points in ZT or CT.

The increased resolution of the STED microscopy did, however, allowed us the estimate the sizes of the PER/BMAL1/CRY1 bodies to be ∼100 nm (Figure 7E and S7D). Because the use of primary and secondary antibodies (size ∼20 nm) for indirect immunolabeling in STED should increase the apparent size of PER bodies, we performed STED microscopy using an anti-GFP nanobody conjugated with STAR 635P (size < 4 nm). As expected, the sizes of PER bodies identified by the GFP nanobody were estimated at 70 nm. The actual sizes of the PER bodies are likely smaller than 70 nm due to the presence of multiple PER2 molecules in unknown orientations in each PER body. This result might be consistent with the 40-nm estimate from the previous single-particle electron microscopy analysis (26). When 40 nm was used as the colocalization distance, the percentage of PER2-EGFP foci that colocalize with BMAL1 foci and CRY1 foci dropped to 8% and 3%, respectively in these 2D image analyses (Figure S7E-F).

To further confirm that most of PER2-EGFP are not associated with CRY1 or BMAL1, we performed immunodepletion assays using protein extracts prepared from the double KI U2OS cells and the lowest concentration of CRY1 or BMAL1 antibodies that could deplete the protein of interest. As shown in Figure 7F, the depletion of CRY1 or BMAL1 did not markedly affect the PER2 levels in the depleted cell extracts. Our conclusion was further confirmed by mPER2 immunodepletion assays by using C57BL/6 mice liver extracts (ZT18, ∼peak of mPER2) (Figure 7G), indicating that our conclusion in U2OS cells can also be applied in mice.

### Most nuclear PER2 and CRY1 do not colocalize in mice SCN cells

Finally, to confirm if our conclusion made from U2OS cells can also be applied in mice SCN cells, we performed Airyscan immunofluorescence imaging experiments to examine the colocalization of PER2 and CRY1 using mice suprachiasmatic nucleus (SCN) tissues. PER2-specific and two different CRY1 antibodies (one mouse antibody and one guinea pig antibody) were used. As expected, mPER2 and mCRY1 foci similar to those in U2OS KI cells were found to be enriched in the nuclei of the SCN cells (Figure 7H and S7G). As expected, despite some colocalization, most mPER2 and mCRY1 foci did not colocalize. In addition, the number of mPER2 foci were significantly higher at ZT12 and CT12 than those at ZT4 and CT4, respectively, consistent with a mPER2 rhythm in SCN cells (Figure 7I). As shown in Figure S7H, the PER2 and CRY1 antibodies had low background nuclear signals in the SCN cells prepared from the *Per1^-/-^Per^2-/-^Per3^-/-^* triple KO and *Cry1*^−/−^*Cry2*^−/−^ double KO mice, respectively (69, 70), indicating the specificity of the antibodies. Together, our results demonstrated that endogenous PER2 only interacts with endogenous BMAL1 and CRY1 transiently and most PER2 bodies are not in complex with BMAL1 or CRY1 at any given time.

## Discussion

LLPS has been used to explain diverse cellular and biochemical processes. The challenges of performing studies of the LLPS behavior of endogenous proteins have meant that many studies were based on protein overexpression in cells or on in vitro biochemical studies (43). Our analysis presents an important cautionary example for LLPS study and demonstrate the importance of examining endogenous proteins in our understanding of biological mechanisms.

PER proteins are the core components of the animal circadian clocks and play essential roles in circadian feedback loops and period length determination (1–3). The PER proteins are IDR-rich and were thought to form super complexes with other clock proteins, prompting us to examine the potential role of LLPS of PER proteins in clock mechanisms. Indeed, even when PER2 was stably overexpressed in U2OS cells at a much lower level than commonly used transient transfection, PER2 LLPS condensates are formed in the nucleus. These condensates satisfied standard LLPS criteria: They were sensitive to concentration changes, fuse with each other, rapidly recover after photobleaching, and are highly sensitive to 1,6-hexanediol treatment. Moreover, the LLPS behavior also appeared to correlate with PER2 biological function as condensate formation depends on PER phosphorylation and the condensates can recruit other clock proteins. Had we relied only on these results, we would have proposed that LLPS was important for clock function. The *Drosophila* PER and the mammalian REV-ERBα were recently shown to exhibit LLPS behavior but such conclusions were also based on protein overexpression in cells (71, 72). How the endogenous dPER and REV-ERBα proteins behave is still unclear.

Super-resolution imaging of the endogenous PER2 protein, however, revealed that the LLPS behavior exhibited by the overexpressed protein was not physiologically relevant. Although the endogenous PER2 formed highly concentrated protein microbodies, these bodies were insensitive to concentration changes, resistant to 1,6-hexanediol treatment, and insensitive to the loss of PER2 phosphorylation. These results indicate that the LLPS of the overexpressed protein and the formation of endogenous PER bodies are mediated by distinct mechanisms. The former is likely mediated by multivalent weak and nonspecific protein interactions involving IDRs but the latter should be caused by strong specific protein-protein interactions insensitive to 1,6-hexanediol treatment and protein phosphorylation. This is different with the protein hub formation of some transcription factors, which was previously shown to be mediated by the multivalent IDR-IDR interactions (43, 46, 47). Although the nature of the protein-protein interactions mediating the PER body formation is not clear, a crystal structure of a mouse PER2 fragment containing the PAS domains revealed a homodimer stabilized by antiparallel packing of the PAS-B β-sheets (73), suggesting that highly specific protein-protein interactions mediated by structured protein regions, rather than unspecific weak multivalent ones, drive the formation of PER bodies (74). Based on our results, we propose that for LLPS studies, endogenous proteins should be examined whenever possible. In addition, the level of the endogenous protein should be compared to that used for LLPS studies. If the former is well below the protein concentration needed for LLPS behavior, LLPS may not be physiologically relevant.

PER proteins play two major roles in the circadian negative feedback loop: They promote the removal of BMAL1-CLOCK complex from E-boxes on chromatin by recruiting CK1 to phosphorylate CLOCK, and they modulate CRY-mediated inhibition of BMAL1-CLOCK activity on chromatin (19–23). PER was thought to mediate these roles through formation of large and stable complexes with CRY1, BMAL1, and CLOCK in the nucleus (26, 28). By directly visualizing the nuclear distribution and dynamic movements of endogenous PER2, BMAL1, and CRY1 using fluorescence reporters, our study provides important insights into the mechanisms of action of these core clock proteins. PER2, BMAL1, and CRY1 all form nuclear microbodies that are under circadian control with PER2 have the most robust rhythm. At their respective peaks, there were about 600 bodies for each of the proteins with PER bodies having the highest amplitude. The number of PER bodies is not solely determined by PER2 levels since at 4 and 10 h after cell synchronization, levels of PER2 were about the same but there were drastically different PER body numbers. The progressive phosphorylation of PER2 during this time frame suggests that phosphorylation may promote PER body formation. However, since mutation of the PER-CK1 interaction domain that abolished PER phosphorylation without significant effects on PER body number or fluorescence intensities in cells, phosphorylation may modulate but is not required for PER body formation.

PER proteins have been previously shown to co-immunoprecipitated with BMAL1 and CRY proteins and have high affinity binding with each other in vitro (2, 4–10), suggesting they can form large stable complex together (26). In contrast, multiple lines of results presented here demonstrate that vast majority of PER microbodies are not in association with BMAL1 or CRY1 bodies at any time, indicating that the interaction between endogenous PER and other clock proteins are transient. These results suggest that the interaction between PER2 and CRY1/BMAL1 in vivo is regulated. Phosphorylation of these proteins is one likely mechanism regulating their interactions.

Live cell imaging of BMAL1-mScarlet-I knock-in cells showed that most BMAL1 bodies move rapidly in the nucleus and that only a small percentage are immobile for up to 16 s, which is similar to observations of other transcription factors (64). Our light-sheet microscopy analyses demonstrated that most PER bodies freely diffuse in the nucleus. Those that are immobile remain so for less than 1 s (average of 0.4 s), suggesting that the association of PER bodies with BMAL1 on DNA is very transient. Thus, PER bodies transiently interact with BMAL1-CLOCK complexes to promote CLOCK phosphorylation by CK1 resulting in dissociation of BMAL1-CLOCK from DNA (19, 20). In live cell imaging of CRY1-mScarlet-I knock-in cells using an Airyscan microscope at 1-s intervals, we did not detect immobile CRY1 bodies, indicating that the association time of CRY1 with chromatin is also much shorter than that of BMAL1. Thus, both PER and CRY1 mediate their roles in the circadian negative feedback process by transient association of with BMAL1 on chromatin. Comparing to the formation of a large and stable clock protein complex, such a property can make PER and CRY1 body to be enzyme-like and act on multiple BMAL1-CLOCK complexes, achieving high efficiency of repression.

The lack of interaction between most PER bodies with CRY bodies was surprising because their levels peak at around the same time during a circadian cycle and the two proteins can be fractionated together (27). In addition, mCRY1 and mPER2 were previously shown to have a high affinity for each other (Kd in the lower nanomolar range) (4), suggesting a stable CRY-PER interaction. Our results here, which are confirmed by fluorescence and immunofluorescence imaging in U2OS cells and SCN cells and immunodepletion assays in cells and liver tissue, and indicate that the PER and CRY interaction is transient in cells, which indicate the existence of a mechanism that can regulate PER-CRY association in vivo. Phosphorylation of either PER or CRY is one likely mechanism that regulates their interaction *in vivo*. CRY can repress BMAL1-CLOCK activity on DNA independently of PER proteins (23, 75, 76). We previously showed that PER and CRY can sustain a functional (but very low amplitude) circadian negative feedback loop by repressing BMAL1-CLOCK activity independently of the PER-CK1 interaction-mediated negative feedback mechanism (19). Although the exact role of PER-CRY association in the clock mechanism is unclear, the transient PER-CRY association may modulate the inhibitory function of CRY to result in rhythmic BMAL1-CLOCK activity. Finally, clock protein overexpression is commonly used in clock studies. As we showed here that the transient transfection- and lentiviral transduction-mediated PER2 expression resulted in over 2,000× and 20× overexpression, respectively. Given the discrepancies between results obtained based on experiments relied on protein overexpression and our findings, our study highlights the importance of studying endogenous proteins to understand clock mechanisms.

Unlike our results here and previous published imaging results of PER2, BMAL1 and CRY1 (29–31, 77), the *Drosophila* PER and CLOCK proteins were recently found to be in a few large discrete foci at the nuclear envelope during the circadian repression phase (78). Such a difference may be caused by differences in circadian clock mechanisms between insects and mammals.

## MATERIALS and METHODS

### Cell Culture Conditions

U2OS, HEK293T and MEF cells were used in this study. HEK293T and MEF cells were cultured in Dulbecco’s Modified Eagle’s Medium (DMEM) with 10% fetal bovine serum (FBS) and penicillin-streptomycin in 35-mm or 10-cm plates. U2OS cells were cultured in McCoy’s 5A medium with 10% FBS and penicillin-streptomycin in 35-mm or 10-cm plates. The cells were maintained in a humidified incubator at 37°C with 5% CO2. All live cell imaging studies were performed at 37°C with CO2. All cell lines generated in this study were derived from U2OS cells, which are originally derived from human bone osteosarcoma epithelial cells of a female patient.

For temperature entrainment, U2OS cells were seeded in 3.5 mm cell culture dishes at 20% confluency, were subjected to a 5-day entrainment with temperature cycles (12 hr at 33°C and 12 hr at 37°C) and then released into constant conditions at 37°C. Medium changes were performed every two days during the transition from 33°C to 37°C. On day 6, cells were harvested and fixed with PFA (paraformaldehyde; Sigma-Aldrich, # 158127) at indicated time points.

### Vector construction, transfection, lentivirus production and stable cell generation

The PER2-EGFP, PER2 (L730G)-EGFP, PER2(1–500)-EGFP, PER2(501–950)-EGFP, and PER2(951–1255) constructs were generated previously (19) and subcloned into the pLX_317 backbone vector using the NEBuilder HiFi cloning kit to generate the lentiviral vectors. The PER2(527-818 S/T-A)-EGFP mutant vector was partially synthesized by Genscript before subcloned into the PER2-EGFP lentiviral vector.

For transfection, Lipofectamine 3000 (Invitrogen) or polyethyleneimine (PEI) transfection reagents were used according to the manufacturer’s instructions. Briefly, cells were seeded in appropriate culture plates and allowed to adhere overnight. The transfection mixture were prepared by diluting the transfection reagents in Opti-MEM medium and adding DNA to the mixture. The mixtures were incubated at room temperature for 20 minutes and then added to the cells. After incubation, the transfection medium was replaced with fresh growth medium, and cells were allowed to recover before further analysis. For the generation of U2OS knock-in and knockout cells, we utilized the Lonza 2D electroporation system (program: X-001) and the Amaxa® Cell Line Nucleofector® Kit V for transfection.

The lentivirus vector and packaging plasmids were co-transfected into HEK293T cells at approximately 70% confluency in a 10 cm dish, and the medium was changed 24 hours later. After 48 hours of incubation, the cell culture supernatant was transferred to a 15 mL centrifuge tube and centrifuged at 3000 RPM for 10 minutes. The supernatant was then filtered through a 0.45 micron syringe filter and collected into a new sterile tube. The viral solution was further concentrated using the Lenti-X™Concentrator (Takara: 631231) according to the manufacturer’s instructions and stored at -80°C for long-term use. The U2OS cells were seeded in a six-well plate and transduced with the lentiviral filtrates 24 hours later in the presence of 8 µg/ml of polybrene. Selection was performed under the pressure of 1 µg/ml of puromycin until control cells completely died. The resultant stably transduced cells were confirmed by confocal microscopy and harvested for further experiments.

### Mice and tissue preparation

C57BL/6J mice were obtained from Jackson Laboratory. Per1-3 triple KO mice and Cry1-2 double KO mice were described previously (69, 70). All animal experiments described in this study were conducted in accordance with the guidelines and approved by the Institutional Animal Care and Use Committee (IACUC) at UT Southwestern Medical Center (APN 2016-101376-G) and were conducted in accordance with the guidelines of the National Institutes of Health Guide for the Care and Use of Laboratory Animals. The animals used in this study were of same genders and within an age range of 8–16 weeks. All mice were housed in pathogen-free barrier facilities under a 12 h/12 h light/dark cycle and at an ambient temperature of 23°C.

All mice were individually housed. After 10 to 12 days of entrainment in 12L:12D, mice were either maintained in LD conditions (ZT 0 is the onset of light) or released in constant darkness (CT condition). At the indicated time points, mice were anesthetized by Ketamine/Xylazine (20 mg/kg Ketamine; 16 mg/kg Xylazine) or at CT4 and CT12 (CT: circadian time. CT12 - onset of activity), followed by immediate cardiac perfused with 0.01M phosphate buffer solution (PBS) and freshly prepared 4 % paraformaldehyde (PFA; Sigma-Aldrich, # 158127) in 0.01M PBS. Anesthesia in DD for CT4 and CT12 was administered under the safe-red light (with Kodak GBX-2 filter) and perfusion was done under the room light. The brains were removed and post-fixed overnight in 4 % PFA in 0.01M PBS. Brains were cryoprotected using 30% sucrose (Crystalgen, # 300-777-1000) solution and embedded in O.C.T. compound (Tissue Plus, Fisher HealthCare, #4585), frozen, and stored at -80 °C until sectioned. 50 mm coronal sections containing the SCN were collected in PBS.

### Western Blot Analysis

The protein concentration of samples was determined by Bradford assay. 50 μg of total protein extracts were separated by SDS-PAGE, transferred onto a PVDF membrane (Millipore), and detected using pierce ECL western blotting substrate (Thermo scientific:32106). The intensities of the bands were quantified using Fiji Image J software. Protein samples with high PER2 levels were diluted to bring relative PER2 levels of different samples to be similar.

### Circadian Bioluminescence Recording

To examine the circadian clock phenotype of the cells of interest, the cells were electroporated with a Per2(E2)-Luc reporter (48) using the Lonza 2D electroporation system (program: X-001) and Amaxa® Cell Line Nucleofector® Kit V. Stable cells were selected with neomycin for 7 days, synchronized with dexamethasone, and bioluminescence was recorded in real-time with the LumiCycle (Actimetrics) at 37°C. The detailed methods for real-time measurement of luminescence were previously described (48, 79).

### Fluorescence recovery after photobleaching

FRAP experiments were conducted with a Zeiss LSM 880 confocal laser scanning microscope equipped with a Plan-Apochromat 63×/1.4 objective lens. Circular regions with diameters ranging from 1 to 3 μm were bleached using a 100% 488 nm/100 mW argon ion laser. The fluorescence signal from the control and bleached area was captured over 70 frames, with acquired images being 512 × 512 pixels, 1.54μs pixel dwell, and 943ms scan time. Fluorescence recovery data were analyzed using FIJI ImageJ software, and the results were averaged using GraphPad Prism.

### Generation of Knock-in cell lines

The knock-in cell lines of PER2-EGFP, BMAL1-mScarlet-I, and CRY1-mScarlet-I were generated using a protocol modified from Gabriel et al (30). To increase knock-in efficiency and eliminate cells with random insertions, we made two main changes. First, we substituted the randomly inserted marker CD4 with mCherry or EYFP. This change allowed us to screen out negative clones with random insertions by using direct fluorescent sorting with FACS. Secondly, to achieve efficient nuclear electroporation with minimal cell death, we used the Lonza™Nucleofector™Transfection 2b Device with nuclear transfection reagents. The nuclear transfection device significantly increased the efficiency of positive clone selection.

The PER2-EGFP donor vector was synthesized by Genscript and subcloned into a CMV-mCherry vector. The three sgRNA plasmids, the PER2-EGFP donor vector, and the i53 plasmid (Addgene:74939) were transfected into U2OS cells by using Lonza 2D system (program: X-001) with Amaxa® Cell Line Nucleofector® Kit V. After 7 days of blasticidin selection, cells were sorted using a BD® LSR II Flow Cytometer and cells that are CFP^+^ mCherry ^−^ were sorted into a mixed population. After 7 days of culture, a CAG-CRE plasmid (Addgene:13775) was transfected into the cells using the Lonza 2D system (program: X-001) with Amaxa® Cell Line Nucleofector® Kit V. After 7 days of blasticidin selection, cells were sorted using a BD® LSR II Flow Cytometer and cells that were CFP+ and mCherry – were sorted into a mixed population. After 7 days of culture, a CAG-CRE plasmid (Addgene:13775) was transfected into the cells using the Lonza 2D system to remove the loxp sites. Afterwards, the cells were sorted into single clones in four 96-well plates. Positive clones were further confirmed by PCR, Western blot, and confocal microscopy. The knock-in cell clones used in this study are homozygous clones.

The knock-in of BMAL1 mScarlet-I and CRY1 mScarlet-I were designed similarly to the PER2-EGFP knock-in, except that the donor vectors were cloned into a CMV-EYFP backbone vector. We created these knock-in cells in the PER2-EGFP KI cell background. Since the PER2-EGFP fluorescence signal is near background level and cannot be detected by FACS, we selected CFP+ and EYFP-cells as candidate double knock-in cell clones. After confirmation by PCR, Western blot, and confocal microscopy, the double KI cells were examined by LumiCycle (LumiCycle, Actimetrics) to detect the presence of normal circadian rhythms. All knock-in cell clones used in this study are homozygous clones.

### Knock-in Point mutation and *Per* gene disruption

The PER2 727-731 in-frame deletion was generated in the PER2-EGFP KI background. Single-stranded DNA (ssDNA) donor with PER2 (VL729/730GG) mutation was synthesized by IDT and co-transfected with sgRNA and Cas9 plasmids using the Lonza 2D system with Amaxa® Cell Line Nucleofector® Kit V. After 7 days of puromycin selection, cells were sorted into single clones in four 96-well plates. The cell clone with in-frame deletion of aa 727-731 were identified using PCR and DNA sequencing: one allele has the in-frame deletion while the other allele has a frameshifting mutation. In addition, we obtained some cell clones with homozygous *Per2* disruption. Disruption of *Per1* gene in the PER2-EGFP KI background was achieved by co-transfecting the two *Per1* sgRNAs and Cas9 plasmids. The resulted clonal cells were screened for clones with both allele of *Per1* gene disrupted by frameshifting mutations.

### Airyscan confocal imaging analyses

Airyscan confocal imaging analyses of cells was performed using a ZEISS LSM880 confocal microscope equipped with Airyscan. Cells were seeded in 35 mm glass bottom dishes (Cellvis: D35-20-1-N), synchronized by adding fresh medium containing 1 µM dexamethasone when cells reached 50% confluency. Cells were fixed with paraformaldehyde at the indicated time points. Nuclei were stained with DAPI. Laser power was adjusted to 0.5-2% following the manufacturer’s instructions. Cells with fewer than 10 passages were used because they exhibit a robust circadian rhythm. To image cellular fluorescence signals, excitation laser wavelengths of 488 nm were used for detecting EGFP while 561 nm was used to detect BMAL1/CRY1-mScarlet. Airyscan detector was employed to detect the signals in super-resolution mode. Imaging was conducted with a 40×/1.3 Plan-Apochromat oil DIC M27 objective, and the pinhole sizes for EGFP and mScarlet were 80 µm, respectively. The frame scan mode was used with a pixel dwell time of 1.19 μs. Filters used were BP420-480+BP495-550, MBS 488/561, and laser power was set to ∼2% for PER2-EGFP knock-in cells. For BMAL1 or CRY1 mScarlet knock-in cells, filters BP570-620+LP645 and MBS 488/561 were used, with laser power set to ∼0.5-2%. Images were averaged 2 times and saved in .czi format, then processed using ZEN 2.3 SP1 software before being converted to .tiff or .ims files for further analyses. Live cell imaging studies were performed at 37°C with CO2.

To distinguish the true knock-in signal from autofluorescence, U2OS cells and PER2 knock-in cells were fixed with paraformaldehyde after 10 hours of dexamethasone synchronization. Lambda scans ((https://www.microscopyu.com/tutorials/lambda-stack-basic-concepts) were performed on a ZEISS LSM880 confocal microscope. The GFP signal was isolated by subtracting its wavelength from all other wavelengths, which were then merged to create a background signal.

Imaris 9.9.1 (Bitplane) software for MAC to track PER2, BMAL1, and CRY1 spots. Time series images of PER2, BMAL1, and CRY1 were captured either by light sheet microscopy (30 ms/frame) or ZEISS LSM880 with Airyscan (0.35-1 s/frame) and exported to Imaris. Spot positions were detected using the Imaris (Bitplane) spot detection function, followed by manual corrections. The tracks were evaluated, and any errors were corrected manually in Imaris.

For 3D imaging, we used a ZEISS LSM880 confocal microscope with airyscan. The obtained images were reconstructed at full resolution. To mark the 3D spots, we utilized IMARIS 9.9.1 (Bitplane) software running on a MAC and created 3D models using the spots rendering method. The grain size for the spots area detail level was set to 0.3 µm, with background subtraction and a filter intensity mean above 11.1. To analyze colocalization, we used Imaris Spot-to-Spot colocalization and colocalization distance analysis, with thresholds set at below 0.1 µm or below 0.04 µm.

For SCN immunofluorescence analyses, SCN slices were washing twice with PBS and permeabilized with 0.2% Tween 20 in PBS for 15 min at room temperature. Next, the slices were blocked with 2% bovine serum albumin (BSA) and 0.1% Tween20 in PBS for 1 h at room temperature. Primary antibodies diluted in the blocking solution and incubated overnight at 4°C (PER2,1:500 PER21-A; CRY1,1:1000# ab54649; CRY1,1:1000# this study). After washing twice with PBS, the slices were incubated with fluorescence-labeled secondary antibodies (1:2000) in the blocking solution for 1 h at room temperature in the dark. The slices were then washed twice with PBS, stained with DAPI (Life Technologies) for 10 min at room temperature in the dark, and washed twice with PBS. Finally, the slices were analyzed using a Zeiss LSM 880 Airyscan confocal microscope.

### Light-Sheet Microscopy of PER2-EGFP KI cells

Cells were imaged with a high-resolution oblique plane light-sheet microscope as described previously (62, 63). Briefly, cells were prepared the same day on 35 mm Mattek dishes with #1.5 coverslips and imaged in in an environment chamber that provided humidity, temperature, and CO2 control. Illumination was provided with 488 and 561 nm lasers, and 525/50 and 593/LP emission filters were used to isolate the fluorescence emission arising from green- and red-emitting fluorophores, respectively, prior to detection with a Hamamatsu Flash 4.0 scientific CMOS camera. Imaging was performed in a 2D format whereby an oblique cross-section of the cell is visualized with a 30-millisecond exposure time for ∼500 timepoints, thereby allowing the observation of rapid PER body dynamics.

### STED microscopy

PER2-EGFP KI cells was synchronized by adding fresh medium containing 1 µM dexamethasone when cells reached 50% confluency. 10 hr after synchronization, the cells were twashed three times with warm PBS and fixed with 4% paraformaldehyde (PFA) for 20 min at 20°C. After washing twice with PBS, cells were permeabilized with 0.2% Tween 20 in PBS for 10 min at room temperature. Next, the cells were blocked with 2% bovine serum albumin (BSA) and 10% (v/v) glycerol in PBS for 1 h at room temperature. Primary antibodies diluted 500-1000 times in the blocking solution were added and incubated overnight at 4°C. After washing twice with PBS, the cells were incubated with fluorescence-labeled secondary antibodies (goat anti-mouse STAR red or goat anti-rabbit STAR orange (1:2000)) in the blocking solution for 1 h at room temperature in the dark. The cells were then washed twice with PBS, stained with DAPI (Life Technologies) for 10 min at room temperature in the dark. For GFP nanobody staining, the primary antibody nano GFP 635P (1:1000 dlution) in the blocking solution were added and incubated overnight at 4°C. The cells were then washed twice with PBS, stained with DAPI (Life Technologies) for 10 min at room temperature in the dark.

The STED imaging was performed with the Facility Line of Abberior Instrument with Olympus 60X oil objective. Single z plain 2D STED image was taken using the Lightbox software. For dual color STED Image, line scan mode was used. STAR RED (Abberior) was imaged with 640 nm excitation laser and 775 nm depletion laser. STAR ORANGE was imaged with 561 excitation laser and 775 nm depletion laser. Line scan mode was used to image STAR RED and STAR ORANGE with parameters, Excitation 640 nm 15% with 30% STED, 775 nm 10% with 20% STED. Dwell Time 5 μs and pixel size 10 nm were used. Images were averaged 5 times. For DAPI, 405 mm excitation with confocal mode was used. All images were taken with 1.0 AU pinhole based on 640 nm.

### Immunofluorescence assays

Cells were washed three times with warm PBS and fixed with 4% PFA for 20 min at 20°C. After washing twice with PBS, cells were permeabilized with 0.2% Tween 20 in PBS for 10 min at room temperature. Next, the cells were blocked with 2% bovine serum albumin (BSA) and 10% (v/v) glycerol in PBS for 1 h at room temperature. Primary antibodies diluted 500-1000 times in the blocking solution were added and incubated overnight at 4°C. After washing twice with PBS, the cells were incubated with fluorescence-labeled secondary antibodies (1:2000) in the blocking solution for 1 h at room temperature in the dark. The cells were then washed twice with PBS, stained with DAPI (Life Technologies) for 10 min at room temperature in the dark, and washed twice with PBS. Finally, the cells were analyzed using a Zeiss LSM 880 confocal microscope.

### CRY1 antibody generation

CRY1 polyclonal antibody was raised against mouse CRY1(496–606) recombinant protein in guinea pigs. CRY1-specific antibodies were purified from the antiserum via negative selection followed by affinity selection using NHS-activated Sepharose 4 Fast Flow resin (Cytiva Cat# 17090601) conjugated to Cry1-/-mouse liver and kidney total protein lysates and mCRY1(496–606) recombinant protein, respectively.

### Immunodepletion assay

PER2-EGFP KI; BMAL1-mScarlet-I KI and PER2-EGFP KI; CRY1-mScarlet-I KI U2OS cells were harvested after a 10-hour DXMS synchronization and used to prepare cell extracts. To prepare mice liver extracts, C57BL/6J mice were maintained under a 12:12 hours light-dark (LD) cycle for 14 days and liver tissue were collected at ZT18. The harvested U2OS cells and liver tissue were lysed using RIPA buffer (containing 50mM Tris•HCl at pH 7.6, 150mM NaCl, 1% NP-40, 0.5% sodium deoxycholate, 0.1% SDS and 1x protease inhibitor). The protein concentration was determined using the Bio-Rad Protein Assay Dye Reagent. Subsequently, 30 µg of protein extract was incubated with 0.04 µg of specific antibodies or control IgG bound to 10 µL of magnetic beads (#ThermoFisher 26162) for 3 hours. The unbound fractions were then analyzed by western blotting for protein of interest.

### Conflict of interest

The authors have no declared conflict interests.

## Acknowledgements

We thank Drs. Katherine Phelps and Marcel Mettlen and the UT Southwestern Quantitative Light Microscopy Core for assistances in imaging analyses and Drs. Michael Rosen, Steven McKnight, Carla Green, Joachim Seeman, Matthew Parker and Xuewu Zhang for discussions and suggestions. We would like to thank the National Institutes of Health (R35GM118118 to Y.L.; U54CA268072, P30CA142543, and RM1GM145399 to KMD; R01NS114527 to S. Y.), the Welch Foundation (I-1560 to Y.L.) and the National Science Foundation (IOS-1931115 to S. Y.) for their generous funding. STED microscopy facility were supported by the Peter O’Donnell Jr. Brain Institute NeuroMicroscopy Core.

## Author contributions

P.X., X.X., and Y.L. designed the study. P.C.X., X.X., C.Y., K.M.D., I. L., S. K. T. T., and S.Y. performed experiments and analyzed data. J. T. and Y. X. provided resources and analyzed data. P.C.X., S.Y., K.M.D. and Y.L. wrote the manuscript.

## Notes

### Competing Interest Statement

The authors have declared no competing interest.

## References

1. J. S. Takahashi, Transcriptional architecture of the mammalian circadian clock. Nat Rev Genet 18, 164–179 (2017).

2. C. L. Partch, C. B. Green, J. S. Takahashi, Molecular architecture of the mammalian circadian clock. Trends Cell Biol 24, 90–99 (2014).

3. M. H. Hastings, A. B. Reddy, E. S. Maywood, A clockwork web: circadian timing in brain and periphery, in health and disease. Nature reviews. Neuroscience 4, 649–661 (2003).

4. I. Schmalen et al., Interaction of circadian clock proteins CRY1 and PER2 is modulated by zinc binding and disulfide bond formation. Cell 157, 1203–1215 (2014).

5. S. N. Nangle et al., Molecular assembly of the period-cryptochrome circadian transcriptional repressor complex. Elife 3, e03674 (2014).

6. G. C. G. Parico et al., The human CRY1 tail controls circadian timing by regulating its association with CLOCK:BMAL1. Proc Natl Acad Sci U S A 117, 27971–27979 (2020).

7. H. Xu et al., Cryptochrome 1 regulates the circadian clock through dynamic interactions with the BMAL1 C terminus. Nat Struct Mol Biol 22, 476–484 (2015).

8. J. L. Fribourgh et al., Dynamics at the serine loop underlie differential affinity of cryptochromes for CLOCK:BMAL1 to control circadian timing. Elife 9 (2020).

9. S. Langmesser, T. Tallone, A. Bordon, S. Rusconi, U. Albrecht, Interaction of circadian clock proteins PER2 and CRY with BMAL1 and CLOCK. BMC Mol Biol 9, 41 (2008).

10. C. Lee, J. P. Etchegaray, F. R. Cagampang, A. S. Loudon, S. M. Reppert, Posttranslational mechanisms regulate the mammalian circadian clock. Cell 107, 855–867 (2001).

11. C. Lee, J. P. Etchegaray, F. R. Cagampang, A. S. Loudon, S. M. Reppert, Posttranslational mechanisms regulate the mammalian circadian clock. Cell 107, 855–867 (2001).

12. K. Vanselow et al., Differential effects of PER2 phosphorylation: molecular basis for the human familial advanced sleep phase syndrome (FASPS). Genes Dev 20, 2660–2672 (2006).

13. Y. Xu et al., Functional consequences of a CKIdelta mutation causing familial advanced sleep phase syndrome. Nature 434, 640–644 (2005).

14. K. L. Toh et al., An hPer2 phosphorylation site mutation in familial advanced sleep phase syndrome. Science 291, 1040–1043 (2001).

15. Y. Xu et al., Modeling of a human circadian mutation yields insights into clock regulation by PER2. Cell 128, 59–70 (2007).

16. M. Zhou, J. K. Kim, G. W. Eng, D. B. Forger, D. M. Virshup, A Period2 Phosphoswitch Regulates and Temperature Compensates Circadian Period. Mol Cell 60, 77–88 (2015).

17. R. Narasimamurthy et al., CK1delta/epsilon protein kinase primes the PER2 circadian phosphoswitch. Proc Natl Acad Sci U S A 115, 5986–5991 (2018).

18. S. Masuda et al., Mutation of a PER2 phosphodegron perturbs the circadian phosphoswitch. Proc Natl Acad Sci U S A 117, 10888–10896 (2020).

19. Y. An et al., Decoupling PER phosphorylation, stability and rhythmic expression from circadian clock function by abolishing PER-CK1 interaction. Nat Commun 13, 3991 (2022).

20. X. Cao, Y. Yang, C. P. Selby, Z. Liu, A. Sancar, Molecular mechanism of the repressive phase of the mammalian circadian clock. Proc Natl Acad Sci U S A 118 (2021).

21. K. Kume et al., mCRY1 and mCRY2 are essential components of the negative limb of the circadian clock feedback loop. Cell 98, 193–205 (1999).

22. E. A. Griffin, Jr., D. Staknis, C. J. Weitz, Light-independent role of CRY1 and CRY2 in the mammalian circadian clock. Science 286, 768–771 (1999).

23. R. Ye et al., Dual modes of CLOCK:BMAL1 inhibition mediated by Cryptochrome and Period proteins in the mammalian circadian clock. Genes Dev 28, 1989–1998 (2014).

24. D. McManus et al., Cryptochrome 1 as a state variable of the circadian clockwork of the suprachiasmatic nucleus: Evidence from translational switching. Proc Natl Acad Sci U S A 119, e2203563119 (2022).

25. J. Fu et al., Codon usage affects the structure and function of the Drosophila circadian clock protein PERIOD. Genes Dev 30, 1761–1775 (2016).

26. R. P. Aryal et al., Macromolecular Assemblies of the Mammalian Circadian Clock. Mol Cell 67, 770–782 e776 (2017).

27. X. Cao, L. Wang, C. P. Selby, L. A. Lindsey-Boltz, A. Sancar, Analysis of mammalian circadian clock protein complexes over a circadian cycle. J Biol Chem 299, 102929 (2023).

28. A. A. Koch et al., Quantification of protein abundance and interaction defines a mechanism for operation of the circadian clock. Elife 11 (2022).

29. N. J. Smyllie et al., Visualizing and Quantifying Intracellular Behavior and Abundance of the Core Circadian Clock Protein PERIOD2. Curr Biol 26, 1880–1886 (2016).

30. C. H. Gabriel et al., Live-cell imaging of circadian clock protein dynamics in CRISPR-generated knock-in cells. Nat Commun 12, 3796 (2021).

31. N. J. Smyllie et al., Cryptochrome proteins regulate the circadian intracellular behavior and localization of PER2 in mouse suprachiasmatic nucleus neurons. Proc Natl Acad Sci U S A 119 (2022).

32. R. Ollinger et al., Dynamics of the circadian clock protein PERIOD2 in living cells. J Cell Sci 127, 4322–4328 (2014).

33. A. A. Hyman, C. A. Weber, F. Julicher, Liquid-liquid phase separation in biology. Annu Rev Cell Dev Biol 30, 39–58 (2014).

34. S. F. Banani, H. O. Lee, A. A. Hyman, M. K. Rosen, Biomolecular condensates: organizers of cellular biochemistry. Nat Rev Mol Cell Biol 18, 285–298 (2017).

35. C. P. Brangwynne, Phase transitions and size scaling of membrane-less organelles. J Cell Biol 203, 875–881 (2013).

36. D. Hnisz, K. Shrinivas, R. A. Young, A. K. Chakraborty, P. A. Sharp, A Phase Separation Model for Transcriptional Control. Cell 169, 13–23 (2017).

37. A. R. Strom et al., Phase separation drives heterochromatin domain formation. Nature 547, 241–245 (2017).

38. M. Du, Z. J. Chen, DNA-induced liquid phase condensation of cGAS activates innate immune signaling. Science 361, 704–709 (2018).

39. P. Li et al., Phase transitions in the assembly of multivalent signalling proteins. Nature 483, 336–340 (2012).

40. M. Kato et al., Cell-free formation of RNA granules: low complexity sequence domains form dynamic fibers within hydrogels. Cell 149, 753–767 (2012).

41. I. Kwon et al., Phosphorylation-regulated binding of RNA polymerase II to fibrous polymers of low-complexity domains. Cell 155, 1049–1060 (2013).

42. S. Alberti, A. Gladfelter, T. Mittag, Considerations and Challenges in Studying Liquid-Liquid Phase Separation and Biomolecular Condensates. Cell 176, 419–434 (2019).

43. D. T. McSwiggen, M. Mir, X. Darzacq, R. Tjian, Evaluating phase separation in live cells: diagnosis, caveats, and functional consequences. Genes Dev 33, 1619–1634 (2019).

44. S. Kroschwald, S. Maharana, A. Simon, Hexanediol: a chemical probe to investigate the material properties of membrane-less compartments. Matters 1–7. (2017).

45. Z. Monahan et al., Phosphorylation of the FUS low-complexity domain disrupts phase separation, aggregation, and toxicity. EMBO J 36, 2951–2967 (2017).

46. S. Chong et al., Tuning levels of low-complexity domain interactions to modulate endogenous oncogenic transcription. Mol Cell 82, 2084–2097 e2085 (2022).

47. S. Chong et al., Imaging dynamic and selective low-complexity domain interactions that control gene transcription. Science 361 (2018).

48. S. H. Yoo et al., A noncanonical E-box enhancer drives mouse Period2 circadian oscillations in vivo. Proc Natl Acad Sci U S A 102, 2608–2613 (2005).

49. R. Chen et al., Rhythmic PER abundance defines a critical nodal point for negative feedback within the circadian clock mechanism. Mol Cell 36, 417–430 (2009).

50. K. Miyazaki, M. Mesaki, N. Ishida, Nuclear entry mechanism of rat PER2 (rPER2): role of rPER2 in nuclear localization of CRY protein. Mol Cell Biol 21, 6651–6659 (2001).

51. K. Yagita et al., Nucleocytoplasmic shuttling and mCRY-dependent inhibition of ubiquitylation of the mPER2 clock protein. EMBO J 21, 1301–1314 (2002).

52. K. Yagita et al., Dimerization and nuclear entry of mPER proteins in mammalian cells. Genes Dev 14, 1353–1363 (2000).

53. D. von Stetten, M. Noirclerc-Savoye, J. Goedhart, T. W. Gadella, Jr., A. Royant, Structure of a fluorescent protein from Aequorea victoria bearing the obligate-monomer mutation A206K. Acta Crystallogr Sect F Struct Biol Cryst Commun 68, 878–882 (2012).

54. K. L. Toh et al., An hPer2 phosphorylation site mutation in familial advanced sleep phase syndrome. Science 291, 1040–1043 (2001).

55. C. Lee, D. R. Weaver, S. M. Reppert, Direct association between mouse PERIOD and CKIepsilon is critical for a functioning circadian clock. Mol. Cell. Biol. 24, 584–594 (2004).

56. S. H. Yoo et al., PERIOD2::LUCIFERASE real-time reporting of circadian dynamics reveals persistent circadian oscillations in mouse peripheral tissues. Proc Natl Acad Sci U S A 101, 5339–5346 (2004).

57. X. Wu, J. A. Hammer, ZEISS Airyscan: Optimizing Usage for Fast, Gentle, Super-Resolution Imaging. Methods Mol Biol 2304, 111–130 (2021).

58. J. Huff, The Airyscan detector from ZEISS: confocal imaging with improved signal-to-noise ratio and super-resolution. Nat Methods 12 (2015).

59. M. K. Bunger et al., Mop3 Is an Essential Component of the Master Circadian Pacemaker in Mammals. Cell 103, 1009–1017 (2000).

60. R. Narumi et al., Mass spectrometry-based absolute quantification reveals rhythmic variation of mouse circadian clock proteins. Proc Natl Acad Sci U S A 113, E3461–3467 (2016).

61. N. Koike et al., Transcriptional architecture and chromatin landscape of the core circadian clock in mammals. Science 338, 349–354 (2012).

62. E. Sapoznik et al., A versatile oblique plane microscope for large-scale and high-resolution imaging of subcellular dynamics. Elife 9 (2020).

63. B. J. Chang et al., Real-time multi-angle projection imaging of biological dynamics. Nat Methods 18, 829–834 (2021).

64. I. Izeddin et al., Single-molecule tracking in live cells reveals distinct target-search strategies of transcription factors in the nucleus. Elife 3 (2014).

65. D. T. McSwiggen et al., Evidence for DNA-mediated nuclear compartmentalization distinct from phase separation. Elife 8 (2019).

66. N. Preitner et al., The orphan nuclear receptor REV-ERBalpha controls circadian transcription within the positive limb of the mammalian circadian oscillator. Cell 110, 251–260 (2002).

67. G. Vicidomini, P. Bianchini, A. Diaspro, STED super-resolved microscopy. Nat Methods 15, 173–182 (2018).

68. S. Berning, K. I. Willig, H. Steffens, P. Dibaj, S. W. Hell, Nanoscopy in a living mouse brain. Science 335, 551 (2012).

69. J. S. Pendergast, G. A. Oda, K. D. Niswender, S. Yamazaki, Period determination in the food-entrainable and methamphetamine-sensitive circadian oscillator(s). Proc Natl Acad Sci U S A 109, 14218–14223 (2012).

70. M. H. Vitaterna et al., Differential regulation of mammalian period genes and circadian rhythmicity by cryptochromes 1 and 2. Proc Natl Acad Sci U S A 96, 12114–12119 (1999).

71. M. Li, S. Li, L. Zhang, Phosphorylation Promotes the Accumulation of PERIOD Protein Foci. Research (Wash D C) 6, 0139 (2023).

72. K. Zhu et al., An intrinsically disordered region controlling condensation of a circadian clock component and rhythmic transcription in the liver. Mol Cell 83, 3457–3469 e3457 (2023).

73. S. Hennig et al., Structural and functional analyses of PAS domain interactions of the clock proteins Drosophila PERIOD and mouse PERIOD2. PLoS Biol 7, e94 (2009).

74. A. Musacchio, On the role of phase separation in the biogenesis of membraneless compartments. EMBO J 41, e109952 (2022).

75. K. Kume et al., mCRY1 and mCRY2 are essential components of the negative limb of the circadian clock feedback loop. Cell 98, 193–205 (1999).

76. Y. Y. Chiou et al., Mammalian Period represses and de-represses transcription by displacing CLOCK-BMAL1 from promoters in a Cryptochrome-dependent manner. Proc Natl Acad Sci U S A 113, E6072–E6079 (2016).

77. N. Yang et al., Quantitative live imaging of Venus::BMAL1 in a mouse model reveals complex dynamics of the master circadian clock regulator. PLoS Genet 16, e1008729 (2020).

78. Y. Xiao, Y. Yuan, M. Jimenez, N. Soni, S. Yadlapalli, Clock proteins regulate spatiotemporal organization of clock genes to control circadian rhythms. Proc Natl Acad Sci U S A 118 (2021).

79. G. Shi et al., Distinct Roles of HDAC3 in the Core Circadian Negative Feedback Loop Are Critical for Clock Function. Cell Rep 14, 823–834 (2016).

