## supplemental Figures for "Mammalian circadian clock proteins form dynamic interacting microbodies distinct from phase separation"

**Supplemental Figures and Supplemental Data files**

**Figure S1**. (A) Design of stable PER2-EGFP expression plasmid by viral transduction. (B) Western blot result showing the PER2-GFP levels in U2OS cells that stably overexpresses PER2-EGFP and in cells transiently transfected with the PER2-EGFP expression plasmid. The protein sample from the transiently transfected cells were analyzed at 20x-150x dilutions. (C) Representative confocal images (left) and statistics (right) of nuclear and cytoplasmic PER2-EGFP percentage at 24, 48, 72, and 98 h after transfection of the PER2-EGFP expression plasmid into U2OS cells. Scale bar: 5 µm. (D) Representative images of U2OS cells that transiently transfected with a plasmid that expresses PER2- EGFP(A206K). The image was taken 48 hrs after transfection. Scale bar: 5 µm. (E) Western blot result showing the PER2-EGFP level in the cells that stably overexpress PER2-EGFP or PER2(L730G)-EGFP. (F) Percentages of cells that stably overexpress PER2-EGFP, PER2(L730G)-EGFP, or PER2(527 818 S/T-A)-EGFP that contain nuclear condensates. Data are presented as mean ±SD, unpaired two-tailed Student's t test, ∗∗∗p < 0.0001.

**Figure S2.** (A) Western blot result showing the PER2-EGFP level in the PER2-EGFP KI and PER2-EGFP KI cells treated by Per2-specific siRNA (si-PER2) and in PER2-EGFP KI cells in which Per2 gene was disrupted by CRSPR/Cas9. Bottom: sequencing result showing the homozygous two-nucleotide deletion in the Per2 gene in the PER2 KO cells. (B) Time course 2D Airyscan confocal imaging results of nuclear GFP signals in PER2-EGFP KI cells after dexamethasone synchronization. (C) Time course 2D Airyscan confocal imaging results of nuclear GFP signals in PER2-EGFP KI cells after synchronization by temperature cycles. (D) Densitometric analyses of Western blot results of PER2-EGFP levels in the nuclear and cytoplasmic fractions of the PER2-EGFP KI cells at hr 4 and hr 10 after dexamethasone synchronization. (E) Western blot results comparing the levels of PER2-GFP in the KI cells to that of defined amount of recombinant GFP protein (2x10-10g). Quantification of western blot analysis and estimation of PER2 molecules per nuclear body. Data are presented as mean ±SEM, n=3.

**Figure S3**. (A) Sum of nuclear fluorescence intensity per cell (left) and mean of fluorescence intensity of nuclear PER2 bodies (right) in the PER2-EGFP KI cells with/without hexanediol (1.5%) treatment by hexanediol. (B) Mean of fluorescence intensity of PER2 bodies in the indicated cells. (C) DNA sequencing results showing the homozygous disruption of the Per1 gene in the PER2-EGFP KI and PER2(727-731D)-EGFP KI cells.

**Figure S4**. (A) (left panels) Airyscan fluorescence imaging results showing the specific BMAL1-mScarlet-I and CRY1-mScarlet-I signals in the indicated double KI cells at hr 10 after dexamethasone synchronization. Scale bar: 5 µm. (Right) Western blot analysis using an RFP antibody to detect the indicated BAML1-mScarlet-1 or CRY1-mScarlet-1 protein in the indicated KI cells. (B-C) Enlarged 2D imaging results in Figure 5B-C showing the colocalization of PER2 with BMAL1/CRY1-mScarlet-I in the double KI cells. Scale bar: 0.5 µm. (D) Airyscan fluorescence imaging results of the indicated double KI cells at hr 32 after dexamethasone synchronization.

**Figure S5**. (A) 3D mask result of PER2-EGFP and BMAL1-mScarlet-I bodies in the PER2-

EGFP and BMAL1-mScarlet-I double KI cells using Imaris software. Colocalization spots are identified based on the colocalization distance <0.1 μm. (B) 3D mask result of PER2-EGFP and CRY1-mScarlet-I bodies in the PER2-EGFP and BMAL1-mScarlet-I double KI cells using Imaris software. Colocalization spots are identified based on the colocalization distance <0.1 μm.

**Figure S6**. (A) Number of PER2-BMAL1 colocalization foci (left) and PER2-CRY1 colocalization foci (right) at different time points after dexamethasone synchronization. Colocalization distance<0.04 μm. Data are presented as mean ±SEM, n=8 cells. (B) Percentage of BMAL1 and PER2 foci that are colocalized with each other in the PER2-EGFP and BMAL1-mScarlet-I double KI cells at different time points after dexamethasone synchronization. Colocalization distance<0.04 μm. Data are presented as mean ±SEM, n=8 cells. (C) Percentage of CRY1 and PER2 foci that are colocalized with each other in the PER2-EGFP and CRY1-mScarlet-I double KI cells at different time points after dexamethasone synchronization. Colocalization distance<0.04 μm. Data are presented as mean Å ± SEM, n=8 cells.

**Figure S7**. (A) Immunostaining (left) and western blot (right) results of the PER2-EGFP KI cells and KI cells treated by Per2-specific siRNA (si-PER2) cell lines showing the specificity of the GFP antibody used. Scale bar: 5 μm. (B) Immunostaining (left) and western blot (right) results of U2OS cells with/without treatment by Cry1-specific siRNA (si-CRY1) showing the specificity of the CRY1 antibody used. Scale bar: 5 μm. (C) Immunostaining (left) and western blot (right) results of BMAL1 in the control MEF and BMAL1 KO MEF cell lines showing the specificity of the BMAL1 antibody used (Ab3350). Scale bar: 5 μm. (D) STED microscopy using BMAL1/CRY1-speific antibody to estimate the sizes of BMAL1 and CRY1 nuclear bodies in the PER2-EGFP KI cells. (E) Percentage of PER2 and BMAL1 bodies that are colocalized with each other at 10 hrs after dexamethasone synchronization. Colocalization distance<0.04μm. Data are presented as mean ± SEM, n=8 cells. (F) Percentage of PER2 and CRY1 bodies that are colocalized with each other at 10 hrs after dexamethasone synchronization. Colocalization distance<0.04μm. Data are presented as mean ± SEM, n=3 cells. (G) Immunofluorecence imaging results showing the nuclear distribution of PER2 and mCRY1 proteins and their co-localization in SCN prepared at the indicated time points. Cells from the core region of SCN were selected here. (H) Immunofluorecence imaging results indicate the lack of mPER2 or mCRY1 nuclear signals in cells of mice SCN tissues (ZT12) of the PER triple KO or CRY double KO mice, respectively. A rabbit polyclonal PER2 antibody and two CRY1 antibody made from mouse and guinea pig were used. Cells from the core region of SCN were selected here.

**Supplemental data files**

Supplemental data file 1: A movie related to the result in Figure 1D showing the fusion of two small condensates into one.

Supplemental data file 2: A movie showing the 3D images related to Figure 3D showing the nuclear PER2 bodies in the PER2-EGFP KI cells.

Supplemental data file 3: A movie showing the live cell fluorescence imaging of the PER2-EGFP KI cells using an Airyscan super-resolution microscope in 0.35 s sampling intervals.

Supplemental data file 4: A movie related to the result in Figure 4A showing the nuclear movements of PER2 bodies captured by light sheet microscopy in 30ms interval.

Supplemental data file 5: A movie showing the 3D fluorescence images related to Figure 6A which PER2 (green) and BMAL1 (red) bodies in the PER2-EGFP and BMAL1-mScarlet-I double KI cells at different time points after synchronization.

Supplemental data file 6: A movie showing the 3D fluorescence images related to Figure 6B which PER2 (green) and CRY1 (red) bodies in the PER2-EGFP and CRY1-mScarlet-I double KI cells at different time points after synchronization.

Supplemental data file 7: A movie showing live cell imaging of the BMAL1-mScarlet-I KI cells using an Airyscan super-resolution microscope in ~1s sampling intervals showing the nuclear movements of BMAL1 bodies.

Supplemental data file 8: A movie showing live cell imaging of the CRY1-mScarlet-I KI cells using an Airyscan super-resolution microscope in ~1s sampling intervals showing the nuclear movements of CRY1 bodies.

**Figure S1**

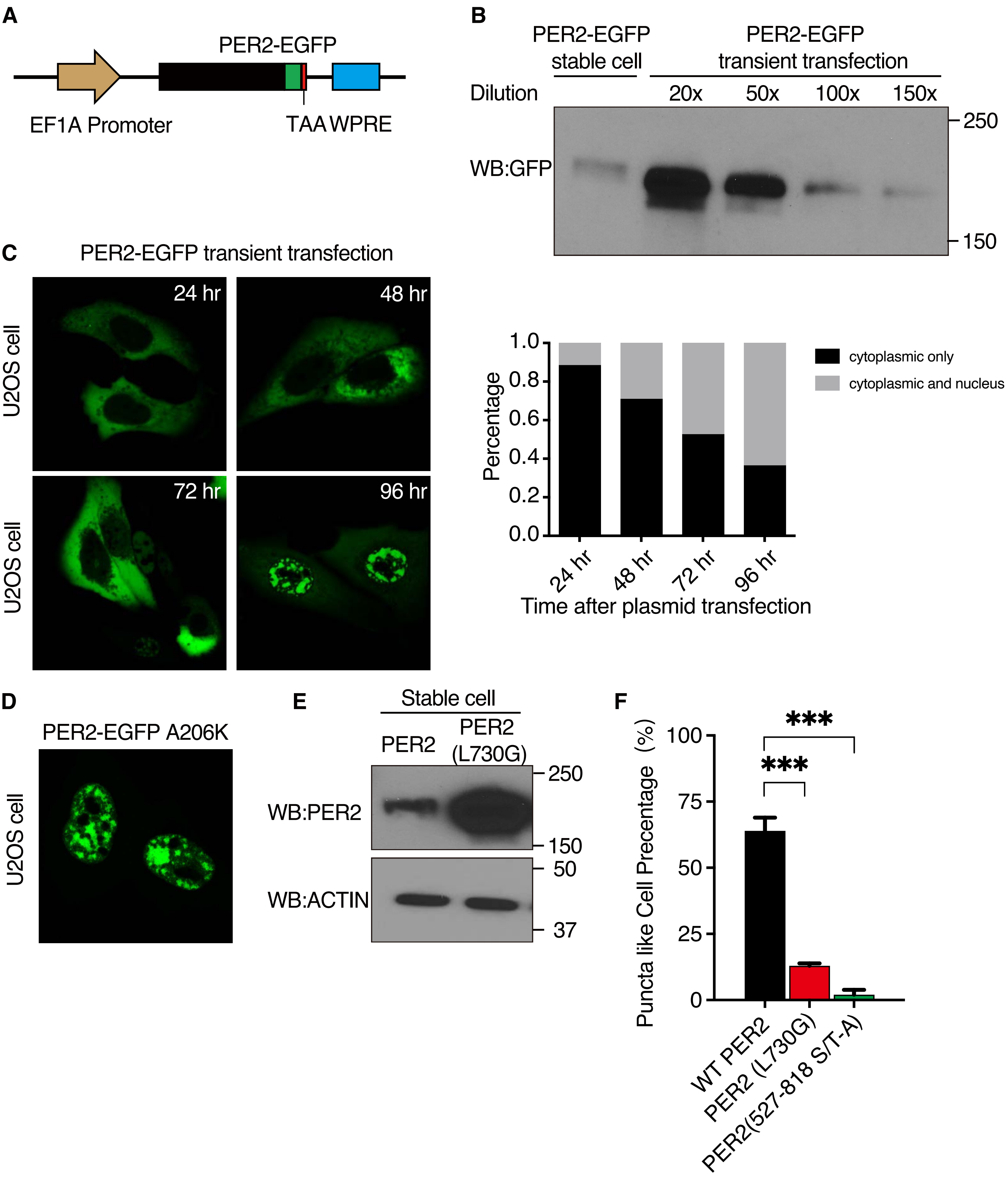

A
PER2-EGFP
EF1A Promoter
TAA
WPRE
B
PER2-EGFP stable cell
PER2-EGFP transient transfection
Dilution
20x
50x
100x
150x
WB:GFP
250
150
C
PER2-EGFP transient transfection
U2OS cell
24 hr
48 hr
72 hr
96 hr
Percentage
cytoplasmic only
cytoplasmic and nucleus
Time after plasmid transfection
D
PER2-EGFP A206K
U2OS cell
E
Stable cell
PER2
PER2 (L730G)
WB:PER2
WB:ACTIN
250
150
50
37
F
Puncta like Cell Precentage (%)
***
***
WT PER2
PER2 (L730G)
PER2(527-818 S/T-A)

**Figure S2**

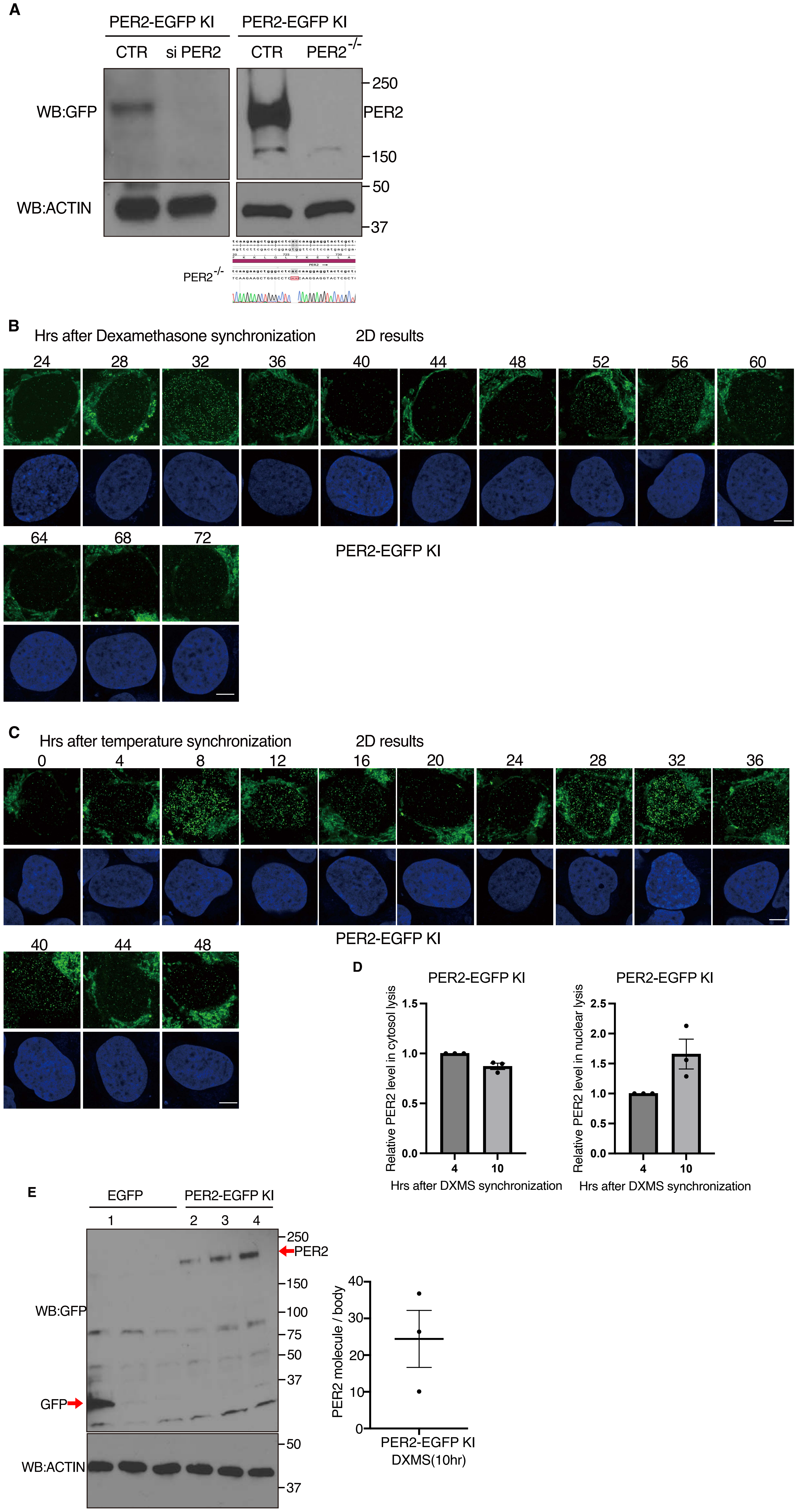
A
PER2-EGFP KI
CTR
si PER2
PER2-EGFP KI
CTR
PER2-/-
WB:GFP
WB:ACTIN
250
PER2
150
50
37
PER2-/-
B
Hrs after Dexamethasone synchronization
2D results
24
28
32
36
40
44
48
52
56
60
64
68
72
PER2-EGFP KI
C
Hrs after temperature synchronization
2D results
0
4
8
12
16
20
24
28
32
36
40
44
48
PER2-EGFP KI
D
PER2-EGFP KI
Relative PER2 level in cytosol lysis
1.5
1.0
0.5
0.0
4
10
Hrs after DXMS synchronization
PER2-EGFP KI
Relative PER2 level in nuclear lysis
2.5
2.0
1.5
1.0
0.5
0.0
4
10
Hrs after DXMS synchronization
E
EGFP
PER2-EGFP KI
1
2
3
4
WB:GFP
GFP
WB:ACTIN
250
PER2
150
100
75
50
37
50
37
PER2 molecule / body
40
30
20
10
0
PER2-EGFP KI
DXMS(10hr)

**Figure S3**

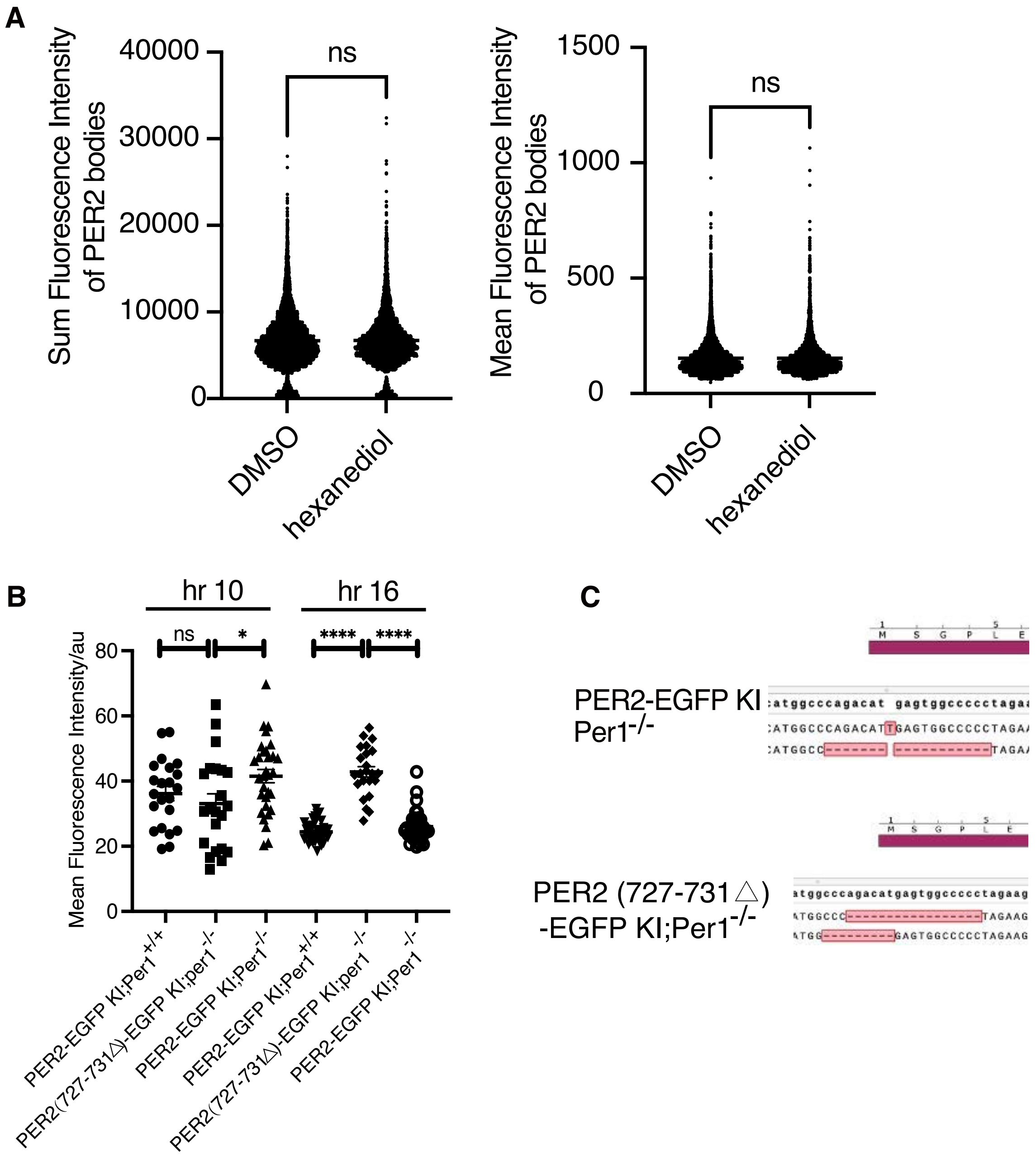
A
Sum Fluorescence Intensity of PER2 bodies
40000
30000
20000
10000
0
ns
DMSO
hexanediol
Mean Fluorescence Intensity of PER2 bodies
1500
1000
500
0
ns
DMSO
hexanediol
B
hr 10
hr 16
ns
*
****
****
Mean Fluorescence Intensity/au
80
60
40
20
0
PER2-EGFP KI;Per1+/+
PER2(727-731△)-EGFP KI;per1-/-
PER2-EGFP KI;Per1-/-
PER2-EGFP KI;Per1+/+
PER2(727-731△)-EGFP KI;per1-/-
PER2-EGFP KI;Per1-/-
C
M S G P L E
PER2-EGFP KI Per1-/-
M S G P L E
PER2 (727-731△) -EGFP KI;Per1-/-

**Figure S4**

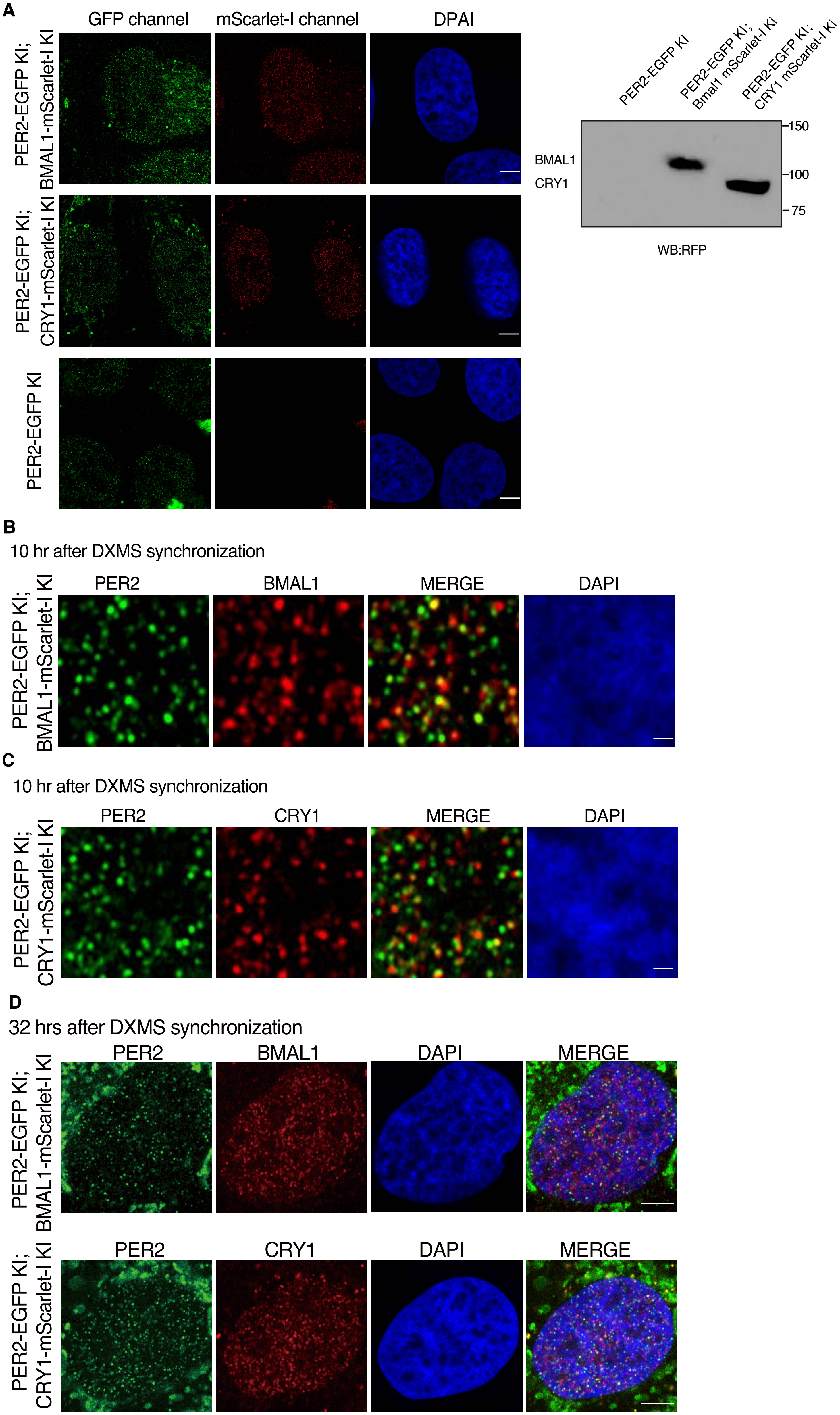
A
GFP channel
mScarlet-I channel
DPAI
PER2-EGFP KI; BMAL1-mScarlet-I KI
PER2-EGFP KI; CRY1-mScarlet-I KI
PER2-EGFP KI
PER2-EGFP KI
PER2-EGFP KI; Bmal1 mScarlet-I Ki
PER2-EGFP KI; CRY1 mScarlet-I Ki
BMAL1
CRY1
150
100
75
WB:RFP
B
10 hr after DXMS synchronization
PER2
BMAL1
MERGE
DAPI
PER2-EGFP KI; BMAL1-mScarlet-I KI
C
10 hr after DXMS synchronization
PER2
CRY1
MERGE
DAPI
PER2-EGFP KI; CRY1-mScarlet-I KI
D
32 hrs after DXMS synchronization
PER2
BMAL1
DAPI
MERGE
PER2-EGFP KI; BMAL1-mScarlet-I KI
PER2
CRY1
DAPI
MERGE
PER2-EGFP KI; CRY1-mScarlet-I KI

**Figure S5**

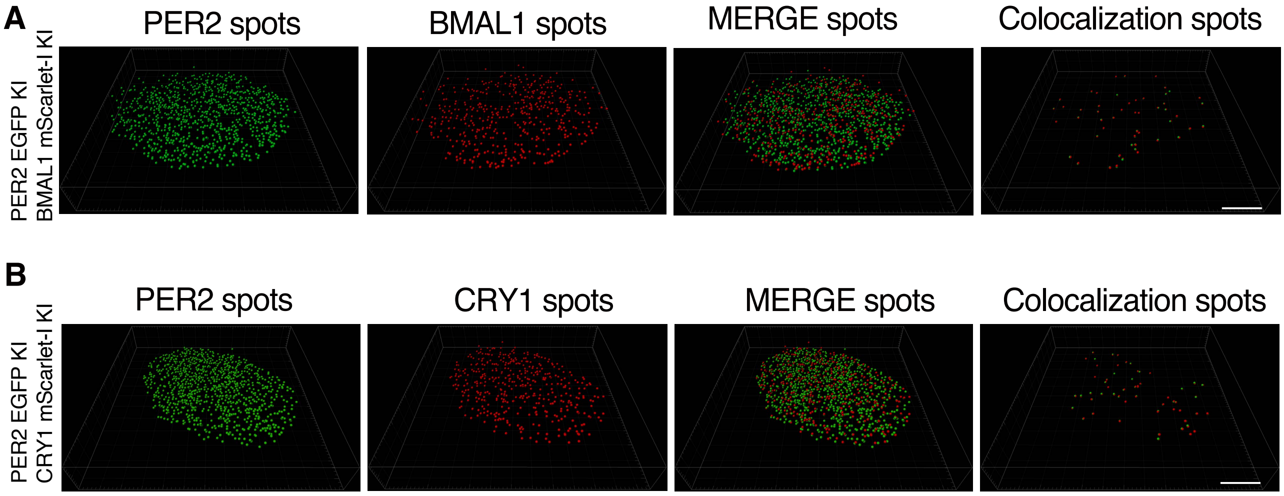
A
PER2 spots
BMAL1 spots
MERGE spots
Colocalization spots
PER2 EGFP KI
BMAL1 mScarlet-I KI
B
PER2 spots
CRY1 spots
MERGE spots
Colocalization spots
PER2 EGFP KI
CRY1 mScarlet-I KI

**Figure S6**

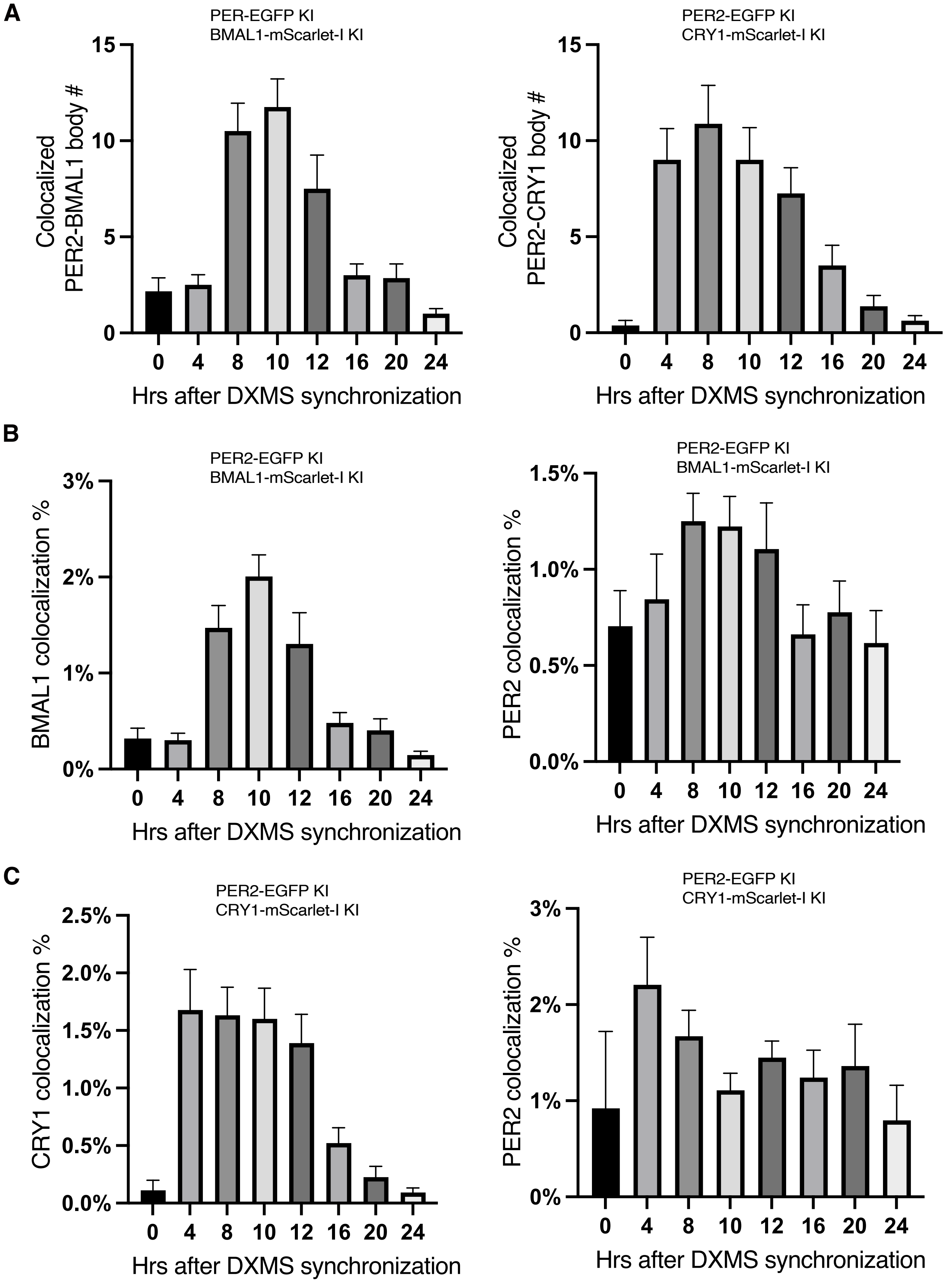
A
PER-EGFP KI
BMAL1-mScarlet-I KI
Colocalized PER2-BMAL1 body #
15
10
5
0
0 4 8 10 12 16 20 24
Hrs after DXMS synchronization
PER2-EGFP KI
CRY1-mScarlet-I KI
Colocalized PER2-CRY1 body #
15
10
5
0
0 4 8 10 12 16 20 24
Hrs after DXMS synchronization
B
PER2-EGFP KI
BMAL1-mScarlet-I KI
BMAL1 colocalization %
3%
2%
1%
0%
0 4 8 10 12 16 20 24
Hrs after DXMS synchronization
PER2-EGFP KI
BMAL1-mScarlet-I KI
PER2 colocalization %
1.5%
1.0%
0.5%
0.0%
0 4 8 10 12 16 20 24
Hrs after DXMS synchronization
C
PER2-EGFP KI
CRY1-mScarlet-I KI
CRY1 colocalization %
2.5%
2.0%
1.5%
1.0%
0.5%
0.0%
0 4 8 10 12 16 20 24
Hrs after DXMS synchronization
PER2-EGFP KI
CRY1-mScarlet-I KI
PER2 colocalization %
3%
2%
1%
0%
0 4 8 10 12 16 20 24
Hrs after DXMS synchronization

Figure S7

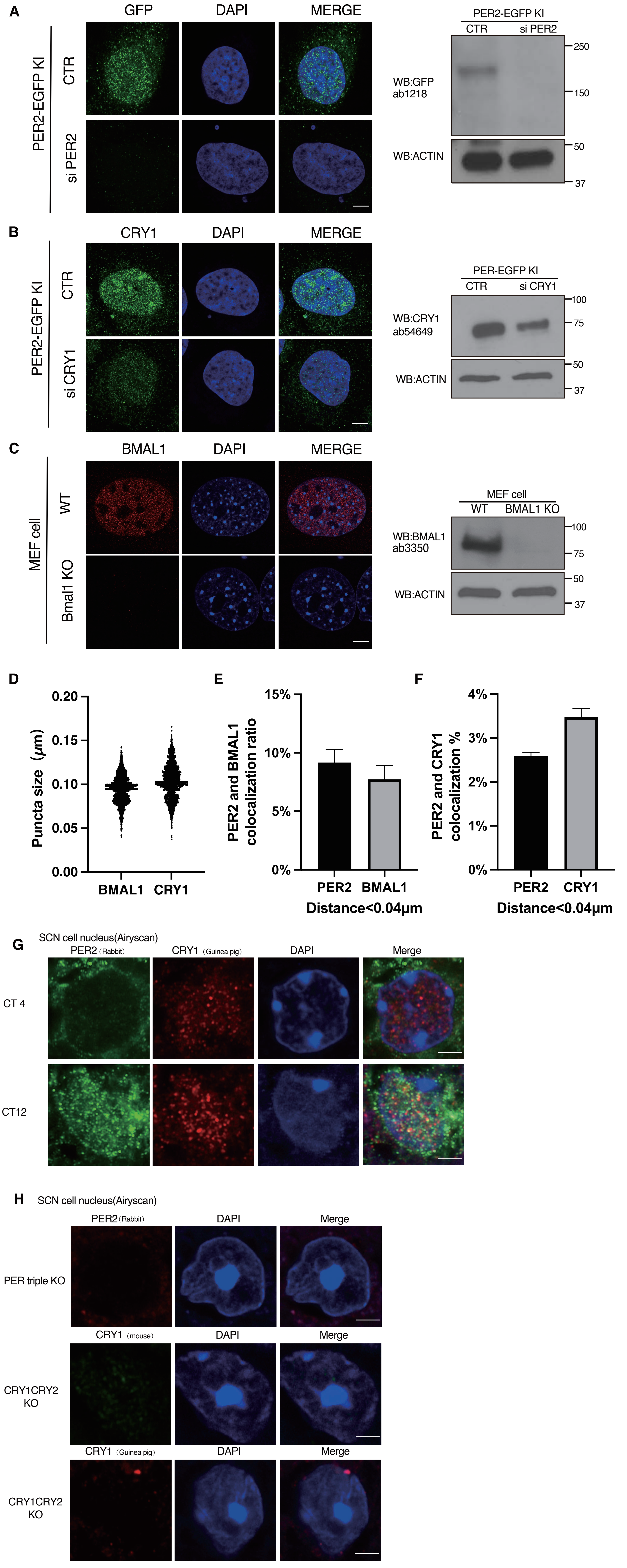
A
GFP
DAPI
MERGE
PER2-EGFP KI
CTR
si PER2
PER2-EGFP KI
CTR
si PER2
WB:GFP
ab1218
WB:ACTIN
250
150
50
37
B
CRY1
DAPI
MERGE
PER2-EGFP KI
CTR
si CRY1
PER-EGFP KI
CTR
si CRY1
WB:CRY1
ab54649
WB:ACTIN
100
75
50
37
C
BMAL1
DAPI
MERGE
MEF cell
WT
Bmal1 KO
MEF cell
WT
BMAL1 KO
WB:BMAL1
ab3350
WB:ACTIN
100
75
50
37
D
Puncta size (μm)
0.20
0.15
0.10
0.05
0.00
BMAL1
CRY1
E
PER2 and BMAL1 colocalization ratio
15%
10%
5%
0%
PER2
BMAL1
Distance<0.04μm
F
PER2 and CRY1 colocalization %
4%
3%
2%
1%
0%
PER2
CRY1
Distance<0.04μm
G
SCN cell nucleus(Airyscan)
PER2 (Rabbit)
CRY1 (Guinea pig)
DAPI
Merge
CT 4
CT12
H
SCN cell nucleus(Airyscan)
PER2 (Rabbit)
DAPI
Merge
PER triple KO
CRY1 (mouse)
DAPI
Merge
CRY1CRY2 KO
CRY1 (Guinea pig)
DAPI
Merge
CRY1CRY2 KO
